# Universal method for the gentle isolation of intact microvessels from frozen tissue: a multiomic investigation into the neurovasculature

**DOI:** 10.1101/2023.05.10.540076

**Authors:** Marina Wakid, Daniel Almeida, Zahia Aouabed, Reza Rahimian, Maria Antonietta Davoli, Volodymyr Yerko, Elena Leonova-Erko, Vincent Richard, René Zahedi, Christoph Borchers, Gustavo Turecki, Naguib Mechawar

## Abstract

The neurovascular unit (NVU), comprised of endothelial cells, pericytes, smooth muscle cells, astrocytic endfeet and microglia together with neurons, is paramount for the proper function of the central nervous system. The NVU gatekeeps blood-brain barrier (BBB) properties which, as a system, experiences impairment in several neurological and psychiatric diseases, and contributes to pathogenesis. To better understand function and dysfunction at the NVU, isolation and characterization of the NVU is needed. Here, we describe a singular, standardized protocol to enrich and isolate microvessels from archived snap-frozen human and frozen mouse cerebral cortex using mechanical homogenization and centrifugation-separation that preserves the structural integrity and multicellular composition of microvessel fragments. For the first time, microvessels are isolated from postmortem vmPFC tissue and are comprehensively investigated using both RNA sequencing and Liquid Chromatography with tandem mass spectrometry (LC-MS-MS). Both the transcriptome and proteome are elucidated and compared, demonstrating that the isolated brain microvessel is a robust model for the NVU and can be used to generate highly informative datasets in both physiological and disease contexts.

## 1. Introduction

To meet the metabolic needs of the 86 billion neurons in the human brain, an elaborate 400 mile-long microvascular network^1, 2^ supplies blood flow to the deep structures of the cerebral hemispheres and gatekeeps blood-brain barrier (BBB) properties. Extensive research efforts have underscored the BBB as a highly selective cellular system. Ultrastructurally, the BBB consists of continuous non-fenestrated brain microvascular endothelial cells (BMECs) that precisely regulate movement between the blood and brain interface through the expression of specialized solute carriers and efflux transporters^3–10^. Attached pericytes and astrocytic end-feet, which are embedded within the vascular basement membrane, together with smooth muscle cells, are the cell types that comprise the neurovascular unit (NVU)^11^ and work in concert to implement coordinated vascular responses to central and peripheral signals. Such responses include the continuous delivery of oxygen and glucose to neurons^12–17^, homeostatic maintenance of the brain^18–22^, the regulation of cerebral blood flow^23–27^ and clearance of interstitial fluid^28–31^. An underappreciated lens through which to investigate disease, NVU dysfunction contributes to cognitive decline and neuronal loss in Alzheimer’s disease-related Aβ and tau pathology^32–37^, traumatic brain injury^38–47^, perivenous myelin lesions observed in multiple sclerosis^48–53^, as well as multiphasic changes to BBB permeability exhibited after stroke^54–58^.

Our understanding of neurovascular development and function has been advanced largely by mouse models. Functional characteristics of the BBB are regulated at the transcriptomic level and, in recent years, different methodologies have been employed to investigate the neurovascular transcriptome. Single-cell sequencing studies have leveraged transgenic-reporter claudin-5-GFP^59, 60^, Tie2–eGFP^61^, and Pdgfrb-eGFP^62^ mouse lines in conjunction with fluorescence-activated cell sorting (FACS) to generate highly informative transcriptomic datasets of mouse BMECs and other vascular cell types, whereas other studies have carried out RNA sequencing of BMECs isolated from Rosa-tdTomato;VE-Cadherin-Cre_ERT2_ mouse models of stroke, MS, TBI and seizure^63^. While past mouse data have provided precious insight in defining core NVU gene expression and underscores the relevance of transcriptomic profiling for better understanding neurovascular function (and dysfunction), recent breakthroughs demonstrate that there are numerous species-specific differences between mouse and human neurovasculature, including solute carrier and efflux transporter expression^63–65^. Such findings reveal the partial utility of animal models for the study of human central nervous system (CNS) disease. Due to scarcity of well-preserved human brain tissue available for research, transcriptomic profiling in human brain samples has been considerably more limited, leaving the investigation of vascular cells neglected in favour of non-vascular cell types, such as neurons and oligodendrocytes. In addition, single-cell or single-nucleus sequencing used to profile expression in *all* cell populations have yielded very low populations of endothelial cells and pericytes from human adult and embryonic cortex samples. Although vessel density, as determined in different brain regions, varies between 361 and 811 vessels/mm^2;66–68^ at an overall endothelial cell density of 4,504 ± 2,666 cells/mm^2;69^, such techniques seem to deplete vascular cells/nuclei for reasons that are not understood and have impeded analysis of human neurovascular transcriptomes^70–74^. Recently, detailed transcriptome-wide atlases of human and mouse brain vascular nuclei in health and disease were generated by two independent groups^64, 65^, both addressing the underrepresentation of vascular cell types and elaborating on species-specific differences in NVU gene expression. Such progress and tools deepen our understanding of human NVU function, yet certain limitations that persist challenge further progress in the field: single-cell and single-nucleus sequencing remains inaccessible to many due to high costs and lack of bioinformatic expertise. Moreover, there is heavy reliance on brain banks for well-characterized frozen human brain tissue, which also requires considerable adaptation of techniques initially optimized for fresh tissue and creates even further disparity in how mouse and human brain tissue are utilized, even with the same experimental question in mind. The biological and bioinformatic biases these experimental decisions create, and their extent, are unknown. Understanding the molecular mechanisms of NVU dysfunction can be achieved by examining gene expression changes in brain microvessels in different disease contexts; and the limited number of existing human neurovascular datasets motivates transcriptomic characterization of more human samples. While it is understood that BMECs perform the BBB function and that other vascular-associated cell types critically regulate that function, the study of microvessels as a preserved unit provides greater insight into the neurovasculature in a manner that dissociated cell types cannot. To this aim, we use a singular, standardized protocol to enrich and isolate microvessels from archived snap-frozen human and frozen mouse cerebral cortex using mechanical homogenization and centrifugation-separation that is gentle enough to dissociate brain tissue while preserving the structural integrity and multicellular composition of microvessel fragments.

The common issue of multiple or contaminating cell types in samples used for tissue-derived RNA sequencing has been largely eliminated by single-cell workflows. However, single-cell and single-nucleus workflows introduce other significant challenges: measurements typically suffer from large fractions of observed zeros possibly due to technical limitations or randomness^75, 76^. Moreover, tissue dissociation and storage biases can induce unwanted transcriptomic alterations and cell type composition differences^77^. Because of this, bulk and single-cell sequencing are complementary strategies in which the former approach warrants a versatile and effective method for isolating the NVU from human brain. Several approaches attempting to investigate the isolated NVU have been developed in the past, albeit with major caveats: FACS-sorting with antibodies against PECAM1/CD31 and CD13 (targeting the endothelial cell membrane and pericyte cell membrane, respectively) require fresh brain tissue^78^, as do other iterations of microvessel isolation for the purpose of cell culture expansion^79, 80^. Past attempts at selective capture of endothelial and vascular-associated cells from frozen human brain have relied exclusively on laser capture microdissection (LCM)^81–84^, which demands considerable optimization if microvessels are to be used for high-throughput applications downstream to laser-capture^85^. To date, microvessels obtained from archived postmortem brain tissue have only been suitable for qPCR and western blot^86^, and more comprehensive knowledge obtained from high-throughput techniques is currently lacking. Critically, microvessels isolated using the described method are in high yield, posses all major vascular-associated cell types, and maintain their *in situ* cellular structure, making them suitable for characterization using high-throughput techniques. The advantages of this simple protocol are manifold: it does not require the experimental setup needed by single-nucleus sorting, nor does it require transgenic mice^87^, enzymatic dissociation^78, 87, 88^, or fresh brain tissue^78, 87, 88^. Importantly, this is the first standardized microvessel isolation method that is demonstrated to work with snap-frozen brain tissue that is compatible across high-throughput downstream applications, removing unknown biases introduced by the use of varied isolation methods.

We have successfully applied the described protocol to postmortem vmPFC tissue from individuals having died suddenly with no neurological nor psychiatric disorder, as well as mouse forebrain tissue as proof of concept that the same procedure can be used in both species. To demonstrate the utility of microvessels isolated from postmortem vmPFC tissue, we processed 5 samples of extracted total RNA and 3 samples of extracted total protein with RNA sequencing and liquid chromatography with tandem mass spectrometry (LC-MS/MS), respectively. Moreover, we sorted BMECs from isolated microvessels using fluorescence-activated nuclei sorting (FANS) as proof of concept that specific neurovascular cell types may be further purified if needed. Bioinformatic processing and analysis of human transcriptomic and proteomic data indicated that isolated samples showed major enrichment for BMEC, pericyte, SMC, and astrocytic endfeet components at both the mRNA and protein level, generating the first multiomic datasets from human brain microvessels.

## 2. Materials and methods

### 2.1. Biological materials

#### Human cortex

This study was approved by the Douglas Hospital Research Ethics Board and written informed consent from next of kin was obtained for each individual included in this study. For each individual, the cause of death was determined by the Quebec Coroner’s office and medical records were obtained. Samples were obtained from Caucasian individuals having died suddenly with no neurological nor psychiatric condition (Table 1). Postmortem brain tissues were provided by the Douglas–Bell Canada Brain Bank (www.douglasbrainbank.ca). Frozen gray matter samples were dissected from the ventromedial prefrontal cortex (vmPFC, Brodmann area 11) by expert brain bank staff stored at –80 °C. The postmortem interval (PMI) is metric for the delay between an individual’s death and collection and processing of the brain. To assess RNA quality, RNA integrity number (RIN) was measured for brain samples using tissue homogenates, with an average value of 5.34. A total of 5 subjects were subjected to RNA-sequencing and 3 subjects were subjects to Liquid Chromatography with tandem mass spectrometry (LC-MS/MS).

**Table 1:** Subject demographics and covariates. Background information on human subjects whose vmPFC tissue was studied. Information includes sex, age, cause of death, postmortem interval (PMI), and tissue pH.

| Experiment type | Brain ID | Sex | Age (years) | Cause of death | PMI (hours) | Tissue pH |
| --- | --- | --- | --- | --- | --- | --- |
| RNA seq | 1 | Female | 76 | Polytrauma (accident) | 26.50 | 6.50 |
| RNA seq | 2 | Female | 51 | Pulmonary embolism | 111.32 | 6.5 |
| RNA seq | 3 | Male | 26 | Polytrauma (car accident) | 12 | 6.75 |
| RNA seq | 4 | Female | 28 | Undetermined | 80 | 6.45 |
| RNA seq | 5 | Female | 45 | Pulmonary embolism | 39.5 | 5.66 |
| Mass spec | 6 | Male | 45 | Gun wound | 20.50 | 6.57 |
| Mass spec | 7 | Male | 54 | Cardiac arrest | 25.25 | 6.61 |
| Mass spec | 8 | Male | 63 | Fall from several metres | 13.00 | 6.84 |

#### Mouse cortex

Male C57BL/6J mice (n = 2) aged between 120-126 days of age were bred, housed and cared for in accordance to the Canadian Council on Animal Care guidelines (CCAC; http://ccac.ca/en_/standards/guidelines), and all methods were approved by the Animal Care Committee from the Douglas Institute Research Center under protocol number 5570. Mice were housed in standard conditions at 22 ± 1 °C with 60% relative humidity, as well as a 12-h light-dark cycle with food and water available *ad libitum*^89^. In accordance with pertinent guidelines and regulations, the mice were anesthetized via intraperitoneal injection of ketamine (10 mg/ml)/xylazine (1 mg/ml) and transcardially perfused with cold PBS 1X. The frontal cortices were removed and immediately frozen in liquid nitrogen and then stored in -80°C freezer.

### 2.2. Tissue homogenization and cellular fractionation

By virtue of a preserved basement membrane, the structure isolated using the described method contains all neurovascular-associated cell types and, therefore, are referred to as “microvessels”. The following protocol describes the specific steps used to isolate and enrich microvessels with retained *in situ* morphology, which is achieved using semi-automated dissociation of microdissected brain tissue into a thorough homogenate followed by low-speed centrifugation (schematic overview in Fig. 1). These steps have been optimized to isolate microvessels from 100 mg of frozen brain tissue, processing a maximum of 4 samples at a time.

**Figure 1.**
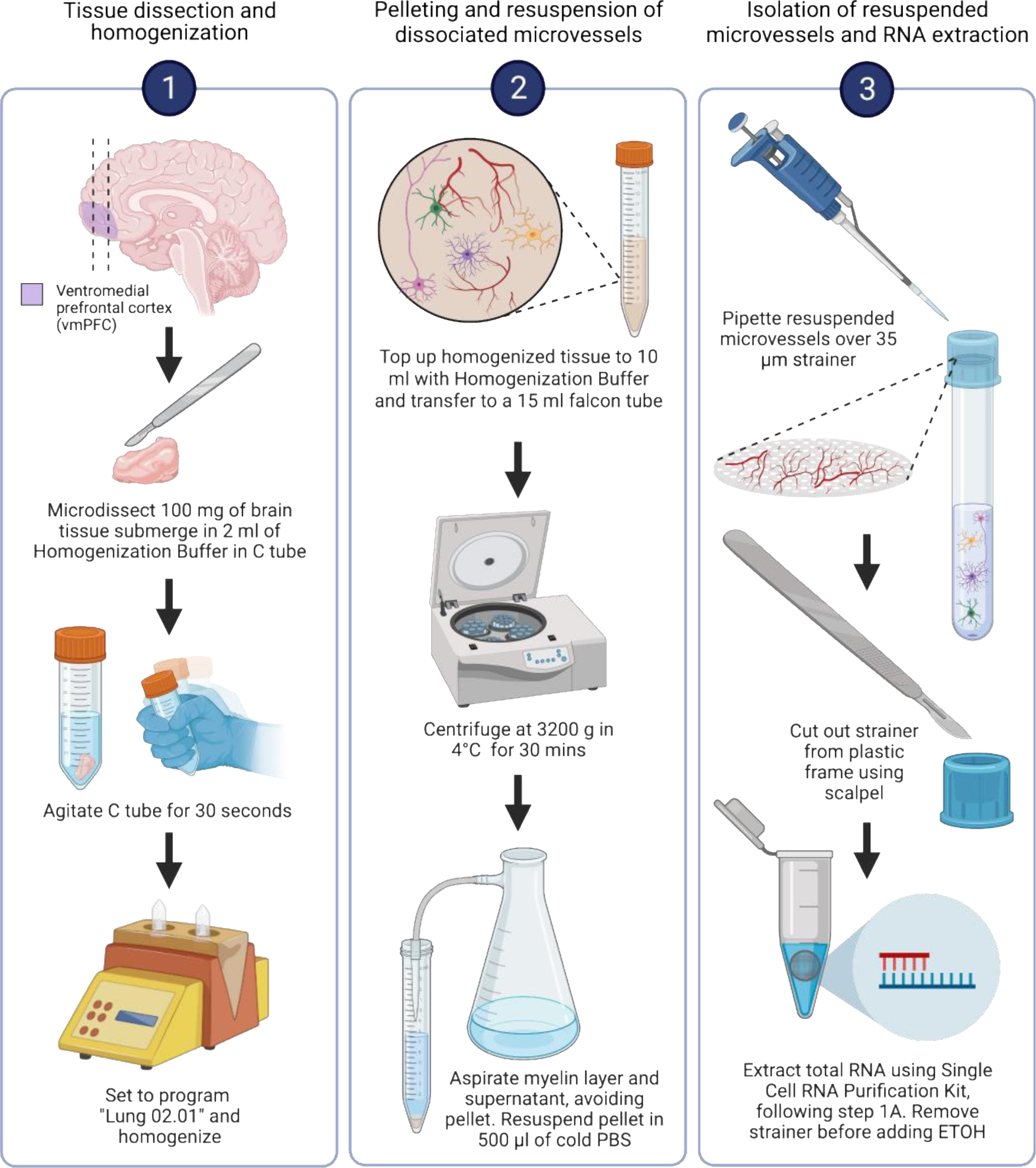
Schematic overview of microvessel isolation from brain tissue samples. Schematic overview of microvessel isolation. After the vmPFC is dissected from coronal slice containing the frontal lobe, 100 mg of vmPFC tissue is more precisely microdissected using a sterile blade or scalpel. Afterwards, the vmPFC tissue sample is dissociated in Homogenization Buffer using the GentleMACS™ Dissociator. Following dissociation, the sample is centrifuged at 3200 g for 30 mins in order to pellet dissociated microvessels. Next, the microvessel-enriched pellet is resuspended in PBS and filtered through a cellular strainer to trap microvessels within the mesh and to allow debris to flow through. The microvessel fragments on top of the mesh can then be subjected to total RNA extraction for downstream, NVU-specific transcriptomic characterization. Additional high-throughput applications can be applied using microvessel-enriched pellets, as discussed in this article. Image generated using BioRender.

Frozen brain tissue was microdissected on top of dry ice using a razor blade, weighed using an analytical scale, and transferred to a chilled 1.5ml Eppendorf tube. The microdissected sample was then transferred to a chilled gentleMACS™ C Tube (Miltenyi Biotec, Bergisch Gladbach, Germany) containing 2 ml of fresh, cold Homogenization Buffer Buffer (1M sucrose + 1% Bovine serum albumin (BSA) dissolved in DEPC-treated water; see supplementary Table 1 for commentary on DEPC-treated water), where the tissue was fully submerged, and the tube was gently agitated for 30 seconds to encourage thawing and osmotic equilibrium (Fig. 2a). The rotating paddle of the GentleMACS™ Dissociator (Miltenyi Biotec, Bergisch Gladbach, Germany) was set to program Lung 02.01 (Fig. 1) and after tissue homogenization was complete, the gentleMACS™ C Tube was returned to wet ice where an additional 8 ml of cold Homogenization Buffer was transferred into the tube, topping up the homogenate to 10 ml, and gently inverted to mix and collect homogenate (Fig. 2b-c). The homogenate was transferred to a chilled 15 ml falcon tube, including any foam produced during paddle rotation (Fig. 2d) and gently inverted a second time in order to prevent the formation of a foamy seal atop of the homogenate when left sitting. The homogenate was centrifuged (Beckman Coulter, model Allegra X-14R) at 3200 g for 30 mins at 4°C. Once centrifugation is complete, a microvessel-enriched pellet forms at the bottom of the falcon tube (Fig. 2e). The supernatant was carefully vacuum-aspirated (which may include an upper layer of clumped, dissociated myelin, similar to milk skin on top of boiled milk) without disturbing the microvessel-enriched pellet (Fig. 2e-f). Anticipated methodological issues are posed in supplementary Table 3, along with potential solutions.

**Figure 2.**
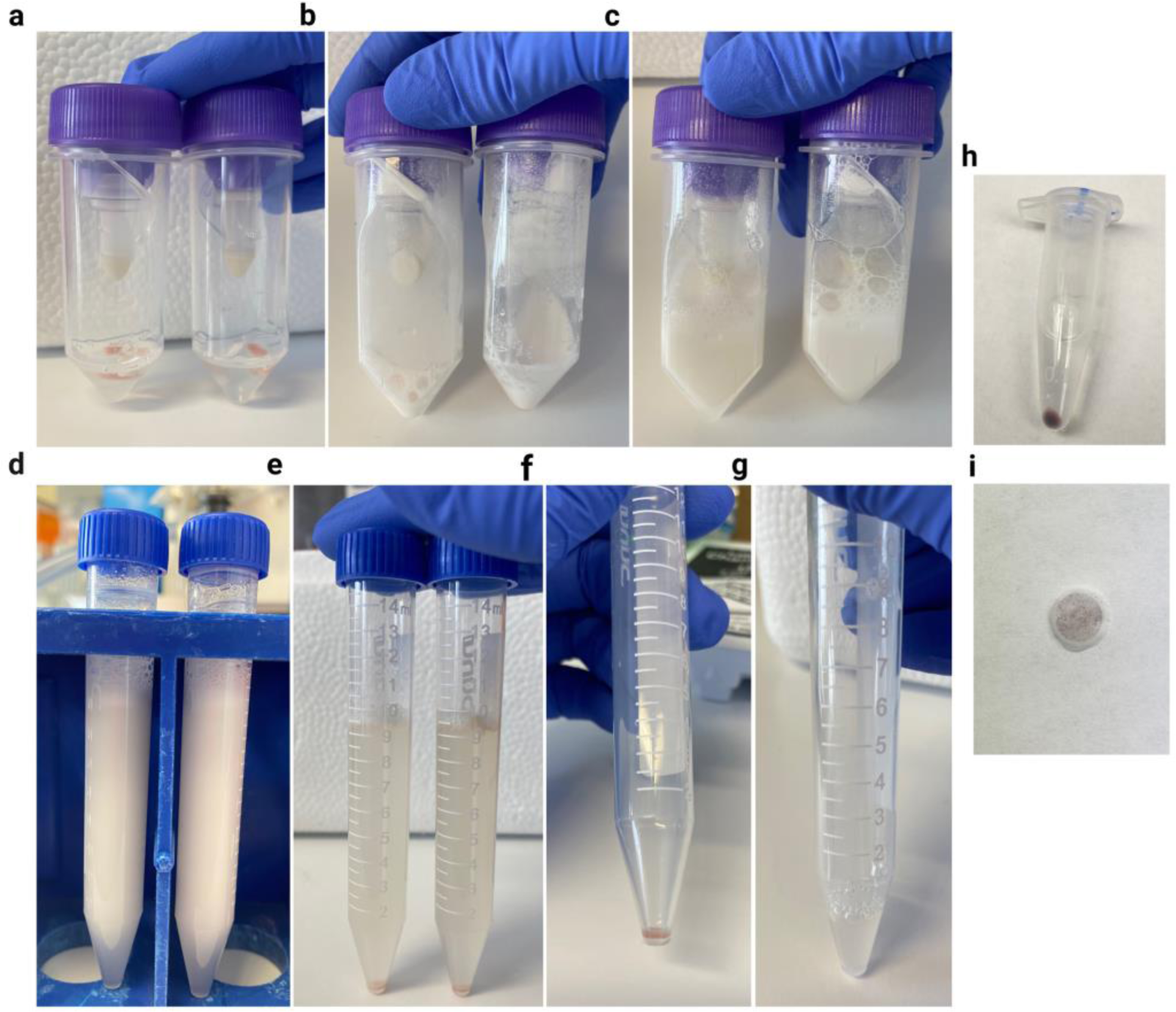
Overview of experimental steps taken to isolate, enrich and collect microvessels. a-g) Overview of experimental steps taken to collect and resuspend pelleted microvessels. a) 100 mg of vmPFC tissue submerged in 2 ml of Homogenization Buffer. b) 100 mg of vmPFC tissue after homogenization. c) Lysate topped-up to 10 ml with Homogenization Buffer. d) Lysate transferred to 15 ml falcon tube. e-f) After centrifugation at 3200 g for 30 min, a microvessel-enriched pellet forms at the bottom of the tube. g) microvessel—enriched pellet is resuspended in 500 ul of PBS. h) Example of BCIP/NBT-stained microvessel pellet, which lends a purple colour to the pellet. i) BCIP/NBT-stained microvessels trapped within the meshwork of a cellular strainer.

### 2.3. Detection of microvessels from the enriched pellet using histology and immunohistochemistry

To first assess yield and stability of microvessel isolation, BCIP/NBT (5-bromo-4-chloro-3’-indolyphosphate and nitro-blue tetrazolium) was used as a chromogenic substrate for endothelial enzyme alkaline phosphatase. Within the brain, alkaline phosphatase activity is localized to cerebral blood vessels^90–96^. BCIP is hydrolyzed by the alkaline phosphatase expressed exclusively in endothelial cells to form a blue intermediate that is then oxidized by NBT to produce a dimer, leaving an intense insoluble purple dye. To this effect, SIGMA*FAST*^™^ BCIP^®^/NBT tablet (Sigma-Aldrich, Missouri, United States) were crushed and then dissolved in the fresh Homogenization Buffer according to manufacturer’s instructions (1 tablet per 10 mls of solution). Crushing the tablet first encourages faster dissolution in what is a highly viscous buffer. Greater immunophenotypic characterization of isolated microvessels was carried out following resuspension of additional microvessel-enriched pellets. Primary antibodies raised against canonical expression markers for BMECs (Vimentin, Laminin and PECAM1), tight junctions (CLDN5), astrocytic end-feet (AQP4), pericytes (PDGFRβ) were utilized, along with appropriate fluorophore-conjugated secondary antibodies to thoroughly characterize collected microvessels. The microvessel-enriched pellet was gently resuspended in 500 μl of cold PBS, with 50 ul transferred into each well of an 8-well chamber slide (Nunc™ Lab-Tek™ II Chamber Slide™ System, Thermo Scientific™, Massachusetts, United States). The chamber slide was then left open faced in a 37°C oven (Fisher Scientific, model Fisherbrand™ Isotemp™) overnight to evaporate the PBS, after which microvessels dry flush to the surface of the slide. It is important to note that immunofluorescent visualization must omit any steps in which the microvessel-enriched pellet is filtered through a cellular strainer because microvessels cannot be released from the strainer and, therefore, cannot be mounted onto a microscope slide. Microvessels were fixed by submerging them to a depth of 2–3 cm with cold 100% methanol for 15 min on ice, after which wells were washed with 1X PBS three times for 5 min. Subsequently, microvessels were incubated in blocking buffer (1% BSA + 0.5% Triton X dissolved in PBS) under agitation for 60 min at 4°C, followed by incubation of each well in 500 ul of primary antibody dilution (1:500 in 1% BSA + 2% normal donkey serum + 0.5% Triton X dissolved in PBS) under agitation overnight at 4°C. Wells were washed with 1X PBS three times for 5 min and finally incubated in fluorophore-conjugated secondary antibody dilution (1:500-1000 in 1% BSA + 2% normal donkey serum + 0.5% Triton X dissolved in PBS) under agitation for 2 hours at room temperature. A final round of washes was followed by removal of the media chamber according to manufacturer’s instructions. Microvessels were coverslipped using VECTASHIELD Antifade Mounting Medium with DAPI (Vector Laboratories, California, United States) and place coverslip.

To expand upon experimental applications possible with isolated human brain microvessels, we adapted several downstream techniques including total RNA extraction, RNA library construction for downstream RNA-sequencing, FANS of BMECs, and total protein extraction for downstream Liquid Chromatography with tandem mass spectrometry (LC-MS/MS).

### 2.4. RNA extraction from isolated microvessels

Total RNA extraction from isolated microvessels is carried out starting with resuspension of the microvessel-enriched pellet. As previously described, the microvessel-enriched pellet was gently resuspended in 500 μl of cold PBS and gradually pipetted through a 35 μm Strainer Cap for FlowTubes™ (Canada Peptide, Quebec, Canada) using vacuum-aspiration underneath to encourage filtration. The result is intact microvessels trapped within the mesh of the strainer, where smaller cellular debris and free-floating nuclei have passed through. After microvessels were trapped within the 35 μm strainer, RNA was extracted using the Single Cell RNA Purification Kit (Norgen Biotek Corp., Ontario, Canada). Using a flat-ended spatula and point-tip forceps, the mesh of the cellular sieve was removed from its plastic frame (Fig. 2i) and immediately submerged into 100 μl of RL buffer in a 1.5 ml Eppendorf tube, according to step 1A of the manufacturer’s protocol. After the addition of 100 μl of fresh 70% ETOH, the solution was pipetted 10 times to wash through the mesh. The mesh was then discarded and the remaining steps according to manufacturer’s instructions were carried out, including the on-column DNase digestion step. Total RNA was quantified using the Agilent TapeStation 2200 (Agilent Technologies, model 2200 TapeStation; for total RNA concentration and RIN, see supplementary Table 2).

### 2.5. Protein extraction from microvessel-enriched pellet or strained microvessels

Total protein has been successfully extracted from either the microvessel-enriched pellet or microvessels stained through a cellular sieve, with one modification to Homogenization Buffer preparation required in which cOmplete^™^ EDTA-free Protease Inhibitor Cocktail (Roche, Basel, Switzerland) and Phosphatase Inhibitor Cocktail 2 (Sigma-Aldrich, Missouri, United States) are added according to the manufacturer’s instructions and based on final volume prepared. With the procedure identical when using strained microvessels (on a sieve), we elaborate protein extraction from frozen microvessel-enriched pellets: Microvessel-enriched pellets were resuspended in 5% sodium dodecyl sulfate (SDS), 100 mM TRIS (pH 7.8) and protein extracted by heating to 99°C for 10 minutes on a hotplate with tube rack. The resuspended pellet was subjected to probe-based sonication (Fisher Scientific, model Thermo Sonic Dismembrator) at 25% amplitude for 3 cycles of 5 seconds and lysates were clarified by centrifugation at 20,000 g for 5 minutes. For estimation of protein concentration, approximately 10% of the sample was aliquoted and diluted to <1% SDS and used for estimation of protein concentration by Pierce™ bicinchoninic acid assay (BCA) Protein Assay Kit (Pierce Biotechnology Inc., Massachusetts, United States). It is important to note that Total protein and total RNA extraction from the same microvessel-enriched pellet has not been validated. Two microvessel-enriched pellets per sample should be prepared, one for RNA and one for protein extraction.

### 2.6. FANS-sorting of BMECs from microvessel-enriched pellet

If further isolation of BMECs from the microvessel enriched pellet is desired, FANS-sorting of ETS-related gene (ERG)^+^ BMECs from the obtained microvessel-enriched pellet can be performed. ERG is a transcription factor whose expression in normal physiological conditions is found exclusively in endothelial nuclei, making it a highly specific pan-endothelial nuclear marker^60, 64, 65, 97, 98^. FANS of BMECs can be carried out following the resuspension of the microvessel-enriched pellet. The pellet is resuspended and centrifuged (Eppendorf, model 5430) at 500 g for 3 min at 4°C, resulting in a pellet once again. The supernatant was aspirated to remove any remaining sucrose, and the washed pellet was resuspended in 250 ul of the following antibody solution: recombinant Alexa Fluor® 647 anti-ERG (1:100 dilution; ab196149, Abcam, Massachusetts, United States) in 0.5% BSA + 0.1% Triton in PBS, incubated under gentle agitation for 2 hours at 4 °C. In the last 10 minutes of the 2-hour incubation, 1 ul of Hoechst 33342 dye (Invitrogen™, Massachusetts, United States) was added to stain nuclear DNA. The suspension was filtered through a 35 μm Strainer Cap for FlowTubes™ and transferred the flow-through into a FACS tube. The BD FACSAria™ Fusion Flow Cytometer (BD Biosciences, California, United States) was used for the sorting of the ERG^+^ population. The gating strategy used for sorting was as follows: doublet discrimination was achieved by the gating of Hoechst 33342 stained singlets in a FSC-A versus Hoechst-A plot using a 350 nm UV laser and a 450/50 filter. The subsequent ERG^+^ population was gated in an Alexa Fluor 647-A vs FSC-A plot using a 640 nm laser in combination with a 730/45 filter. Gating was applied to filter singlets using physical parameters and violet fluorescence (405-nm laser, 525/50 filter). Nonoverlapping gates were adjusted to collect endothelial nuclei based on Alexa Fluor^®^ 647 anti-ERG immunoreactivity (640-nm laser, 730/45 filter). This approach was chosen due to the well-known observation that forward scatter is proportional to size.

### 2.7. Library construction and RNA-sequencing

Microvessel-enriched pellets yielded an average of 8.54 ug/ul of total RNA per sample (supplementary Table 2). Libraries were then constructed using the SMARTer Stranded Total RNA-Seq Kit v3 - Pico Input Mammalian (Takara Bio Inc., Shiga, Japan), which features integration of unique molecular identifiers (UMIs). Libraries were constructed using 10 ng of RNA as input, 2 minutes of fragmentation at 94°C (Applied Biosystems Corporation, model ProFlex PCR), 5 cycles of amplification at PCR1 (addition of Illumina adapters and indexes), 12 cycles of amplification at PCR2 (final RNA-seq library amplification) and clean-up of final library using NucleoMag NGS Clean-up and Size Select beads (Takara Bio Inc., Shiga, Japan). Libraries were then quantified at the Genome Quebec Innovation Centre (Montreal, Quebec) using a KAPA Library Quantification kit (Kapa Biosystems, USA), and average fragment size was determined using a LabChip GX (PerkinElmer, USA) instrument. Libraries were sequenced on the NovaSeq 6000 system (Illumina, Inc., California, United States) using S4 flow cells with 100bp PE sequencing kits (schematic overview in Fig. 3).

**Figure 3.**
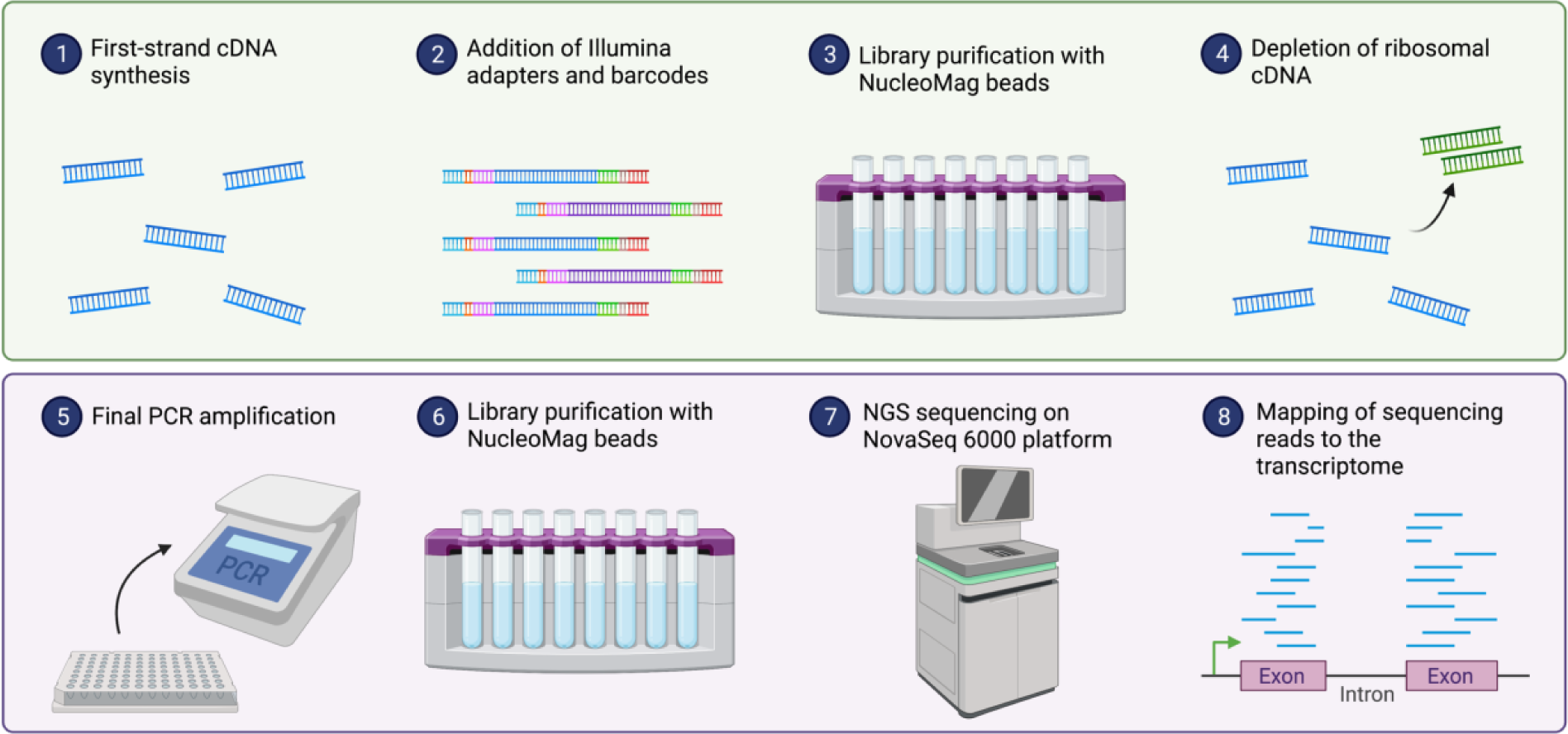
Overview of library preparation and construction for subsequent RNA-sequencing using total RNA extracted from isolated microvessels. Overview of library preparation and construction for subsequent RNA-sequencing using total RNA extracted from isolated microvessels. Image generated using BioRender.

### 2.8. Bioinformatic pipeline and analyses of RNA sequencing data UMI extraction, alignment, de-duplication, metrics and gene counting

RNA sequencing of microvessel libraries yielded an average ∼72 million reads per library, which were then processed following our in-house bioinformatic pipeline. Briefly, UMI extraction based on fastq files was performed using the module extract of umi_tools (v.1.1.2)^99^. Reads were then aligned to the Human Reference Genome (GRCh38) using the STAR software v2.5.4b^100^ with Ensembl v90 as the annotation file and using the parameters: **--twopassMode Basic --outSAMprimaryFlag AllBestScore --outFilterIntronMotifs RemoveNoncanonical --outSAMtype BAM SortedByCoordinate --quantMode TranscriptomeSAM GeneCounts**. Resultant bam files were then sorted and indexed using SAMtools (v.1.3.1)^101^ and duplicate reads with the same UMI were removed using dedup module of umi_tools (v.1.1.2)^99^. Different metrics including the fraction of the exonic, intronic and intergenic reads were calculated using CollectRnaSeqMetrics module of Picard (version 1.129; Picard2019toolkit, Broad Institute, GitHub repository). The expected counts and the transcripts per million (TPMs) were generated using RSEM (v1.3.3; reverse strand mode)^102^.

### Computational deconvolution of RNA sequencing data

Two approaches were used for computational deconvolution of RNA sequencing data. The first approach was performed using the web tool BrainDeconvShiny (https://voineagulab.shinyapps.io/BrainDeconvShiny/), which implements the best performing algorithms and all cell-type signatures for brain, as well as goodness-of-fit calculations based on benchmark work conducted by Sutton et al. (2022)^103^. UMI counts for each gene were converted to transcripts per million (TPMs) to account for the varying length of gene and sequencing depth of each sample, facilitating comparisons across samples. Genes with zero TPMs were then removed, 34370 genes from the original 58303 passed this QC criteria and were used as input into the BrainDeconvShiny tool. Deconvolution was performed twice: the first approach used average expression in control samples from the Velmeshev et al. (2019)^71^ single nuclei dataset (raw data available through the Sequence Read Archive, accession number PRJNA434002; analyzed data available at https://autism.cells.ucsc.edu) as the reference signature for annotated cell types and CIBERSORT v1.04 algorithm to deconvolute sample profiles and estimate cell-type composition. The second approach used the MultiBrain (MB) composite signature^103^ generated by averaging the expression signatures of five datasets for five cell types (neurons, astrocytes, oligodendrocytes, microglia, and endothelia). MB was used as cell type signatures for deconvolution of our dataset using CIBERSORT v1.04.

The second approach to deconvolution was performed in-house. Postmortem NVU single-nucleus data generated on the 10X Genomics Chromium system was accessed from Yang et al. (2022; raw sequencing data are accessible on GEO using the accession code GSE163577)^65^ and used as the reference signature. Seurat^104^ was used to pre-process raw count expression data, removing genes with less than 3 cells or cells with less than 200 expressed genes. 23054 genes from a total of 23537 and 141468 nuclei from a total of 143793 passed these QC criteria. To generate the Yang signature input for the CIBERSORTX tool^105^, counts per million (CPM) values were averaged across nuclei of each cell type.

### 2.9. Gas Chromatography−Mass Spectrometry (LC-MS) with tandem mass tag fractionation

#### Protein digestion and TMT labelling

Extracted proteins were reduced with 20 mM tris(2-carboxyethyl)phosphine (TCEP) at 60°C prior to alkylation with 25 mM iodoacetamide at room temperature for 30 minutes in the dark. An equivalent of 10 μg of total protein was used for proteolytic digestion using suspension trapping (S-TRAP). Briefly, samples were acidified with phosphoric (1.3% final concentration), and then diluted 6-fold in STRAP loading buffer (9:1 methanol:water in 100 mM TEAB, pH 8.5). Samples were loaded onto S-TRAP Micro cartridges (Protifi LLC, Huntington, NY) prior to centrifugation at 2000 g for 2 minutes and washed three times with 50 μl of STRAP loading buffer. Proteins were digested with trypsin (Sigma Corporation, Kanagawa, Japan) at a 1:10 enzyme to substrate ratio for 2 hours at 47°C. Peptides were sequentially eluted in 100 mM TEAB, 0.1% formic acid in water, and 50% acetonitrile, and lyophilized to dryness prior to labelling with TMT 10plex reagents according to the vendor’s specifications (Thermo Fisher Scientific, Massachusetts, United States).

#### Offline high-pH reversed-phase fractionation

Labelled peptides were pooled and again lyophilized to dryness, and then reconstituted in 5 mM ammonium formate and fractionated offline by high pH reversed-phase separation using a Waters Xbridge Peptide BEH C18 column (2.1 x 150mm, 2.5 μm) (Waters Corp., Massachusetts, United States) and an Agilent 1290 LC system (Agilent Technologies, California, United States). Binary gradient elution was performed at a flow rate of 400 μL/minute using mobile phase A) 5 mM ammonium formate adjusted with ammonium hydroxide to pH 10, and B) 100% acetonitrile using the following program: 0 min, 0% B ; 2min, 0% B ; 2.1min, 5% B ; 25min, 30% B ; 30min, 80% B ; 32min, 80% B ; 2 min post-run, 0% B. Fractions were collected every 30 seconds and the first and last 7 fractions were concatenated such that even and odd samples were pooled separately, resulting in 20 fractions in total.

#### LC-MS/MS

Samples were analyzed by data dependent acquisition (DDA) using an Easy-nLC 1200 online coupled to a Q Exactive Plus (both Thermo Fisher Scientific, Massachusetts, United States). Samples were first loaded onto a pre-column (Acclaim PepMap 100 C18, 3 μm particle size, 75 μm inner diameter x 2 cm length) in 0.1% formic acid (buffer A). Peptides were then separated using a 60-min binary gradient ranging from 3-40% B (84% acetonitrile, 0.1% formic acid) on the analytical column (Acclaim PepMap 100 C18, 2 μm particle size, 75 μm inner diameter x 25 cm length) at 300 nL/min. MS spectra were acquired from m/z 350-1,500 at a resolution of 70,000, with an automatic gain control (AGC) target of 1 x 10^6^ ions and a maximum injection time of 50 ms. The 15 most intense ions (charge states +2 to +4) were isolated with a window of m/z 1.2, an AGC target of 2 x 104 and a maximum injection time of 64 ms and fragmented using a normalized higher-energy collisional dissociation (HCD) energy of 28. MS/MS spectra were acquired at a resolution of 17,500 and the dynamic exclusion was set to 30 s. DDA MS raw data was processed with Proteome Discoverer 2.5 (Thermo Scientific, Massachusetts, United States) and searched using Sequest HT against a FASTA file containing all reviewed protein sequences of the canonical human proteome without isoforms downloaded from Uniprot (https://www.uniprot.org). The enzyme specificity was set to trypsin with a maximum of 2 missed cleavages. Carbamidomethylation of cysteine was set as static modification and methionine oxidation as variable modification. The precursor ion mass tolerance was set to 10 ppm, and the product ion mass tolerance was set to 0.02 Da. The percolator node was used and the data was filtered using a false discovery rate (FDR) cut-off of 1% at both the peptide and protein level. The Minora feature detector node of Proteome Discoverer was used for precursor-based label free quantitation.

## 3. Results

### 3.1. Immunophenotypic characterization reveals isolated brain microvessels have preserved morphology and expression integrity

Human brain microvessels were isolated from 5 frozen vmPFC grey matter samples microdissected from healthy individuals who died from peripheral diseases or natural events (Table 1). Because the same basement membrane that maintains the integrity of the endothelium also ensheaths astrocytic endfeet as well as pericytes or smooth muscle cells, it is impossible to isolate solely BMECs and, therefore, isolated microvessels also contained microvessel-associated cell types. Importantly, the described method allows for isolation of microvessels from other brain regions (data not shown) and has additionally been performed in the dorsolateral prefrontal cortex (dlPFC, Brodmann area 8/9), the primary visual cortex (Brodmann area 17), as well as the hippocampus (Brodmann area 28).

Following isolation of human brain microvessels, chromogenic staining of resuspended pellets using BCIP/NBT (5-bromo-4-chloro-3’-indolyphosphate and nitro-blue tetrazolium), a substrate for endothelial enzyme alkaline phosphatase, demonstrated isolation and enrichment of predominantly microvessels from vmPFC tissue samples (Fig. 4a-b), with similar success when isolating microvessels from mouse cortex (Fig. 4c). The cytoarchitecture of brain microvessels is both complex and comprised of several cell types, thus, following isolation of brain microvessels, we aimed to characterize the structure and morphological integrity of isolated microvessels. Immunophenotypic characterization of several NVU markers revealed expression of vimentin (VIM; anti-Vimentin antibody RV202, Abcam), laminins (LAM; anti-Laminin antibody L9393, Sigma-Aldrich), claudin 5 (CLDN5, anti-Claudin 5 antibody ab15106, Abcam), platelet derived growth factor receptor beta (PDGFRβ, anti-PDGFRβ monoclonal antibody G.290.3, Thermo Fisher Scientific) and aquaporin 4 (AQP4, anti-Aquaporin 4 antibody [4/18], Abcam). More precisely, vimentin (Fig. 5a-b), a regulator of actin cytoskeleton in smooth muscle^106^, and laminins (Fig. 5c), the major basement membrane component responsible for signal transduction via interaction with cell surface receptors^107^, are expressed continuously and homogenously across the entire length of the endothelial surface. CLDN5, a major functional constituent of tight junctions^108^, was also stained with no apparent discontinuity, suggesting that endothelial tight junctions were preserved (Fig. 5d). Pericyte coverage of BMECs, immunolabeled by regulator of angiogenesis and vascular stability PDGFRβ^109^ was detected adhering to the surface of microvessels (Fig. 5e). These results suggest that the overall *in situ* brain microvessel structure is conserved after isolation.

**Figure 4.**
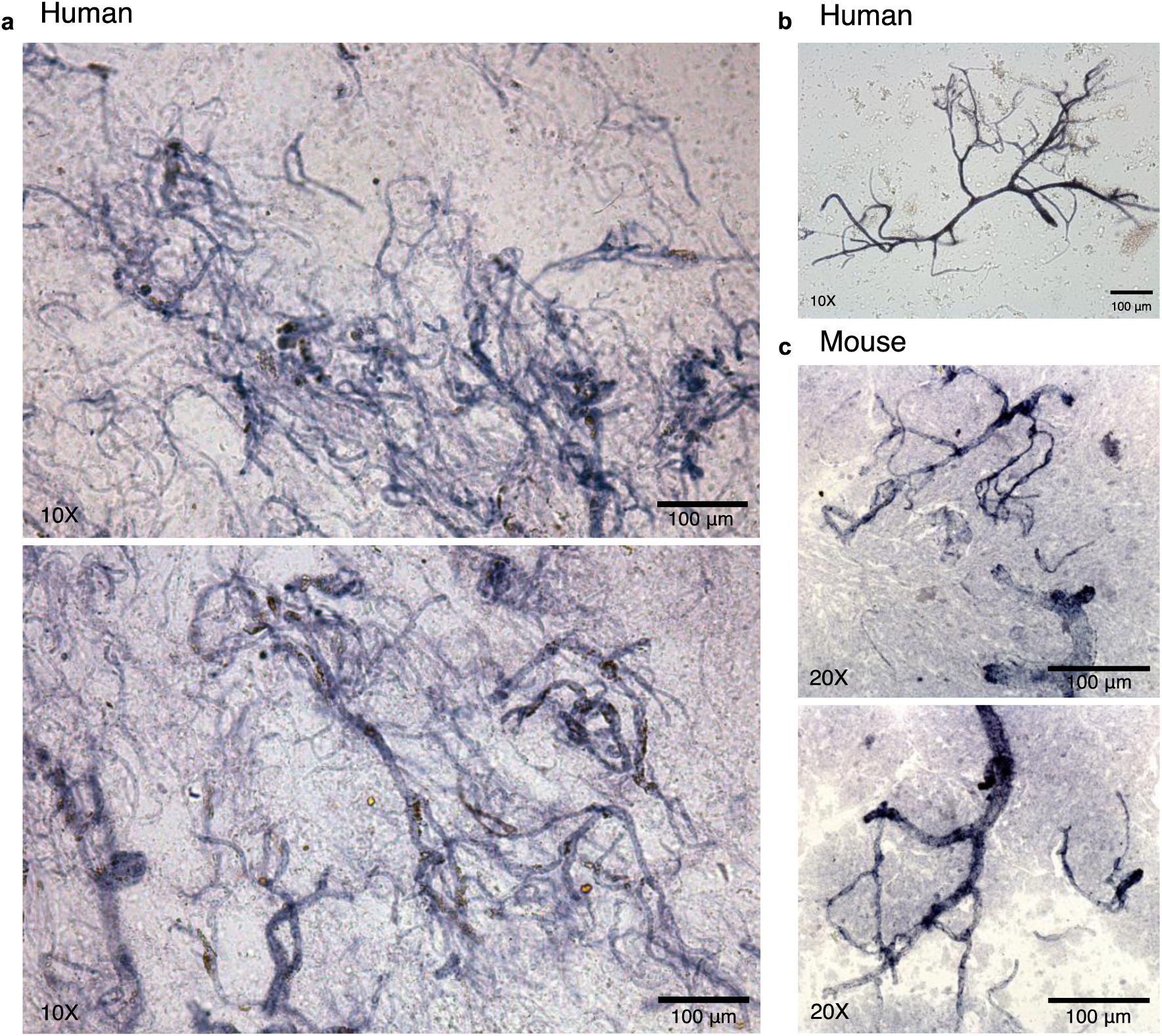
Isolated microvessels are enriched from postmortem vmPFC tissue following the described protocol. a-c) Isolated microvessels are enriched from brain tissue following the described protocol. a-c) Brightfield images of chromogenically stained microvessels using BCIP/NBT substrate demonstrated isolation and enrichment of predominantly microvessels from postmortem vmPFC tissue samples. b) example of preserved microvessel morphology and integrity isolated from postmortem vmPFC tissue. c) Similar microvessel enrichment was observed when the same protocol was carried out using mouse cortex.

**Figure 5.**
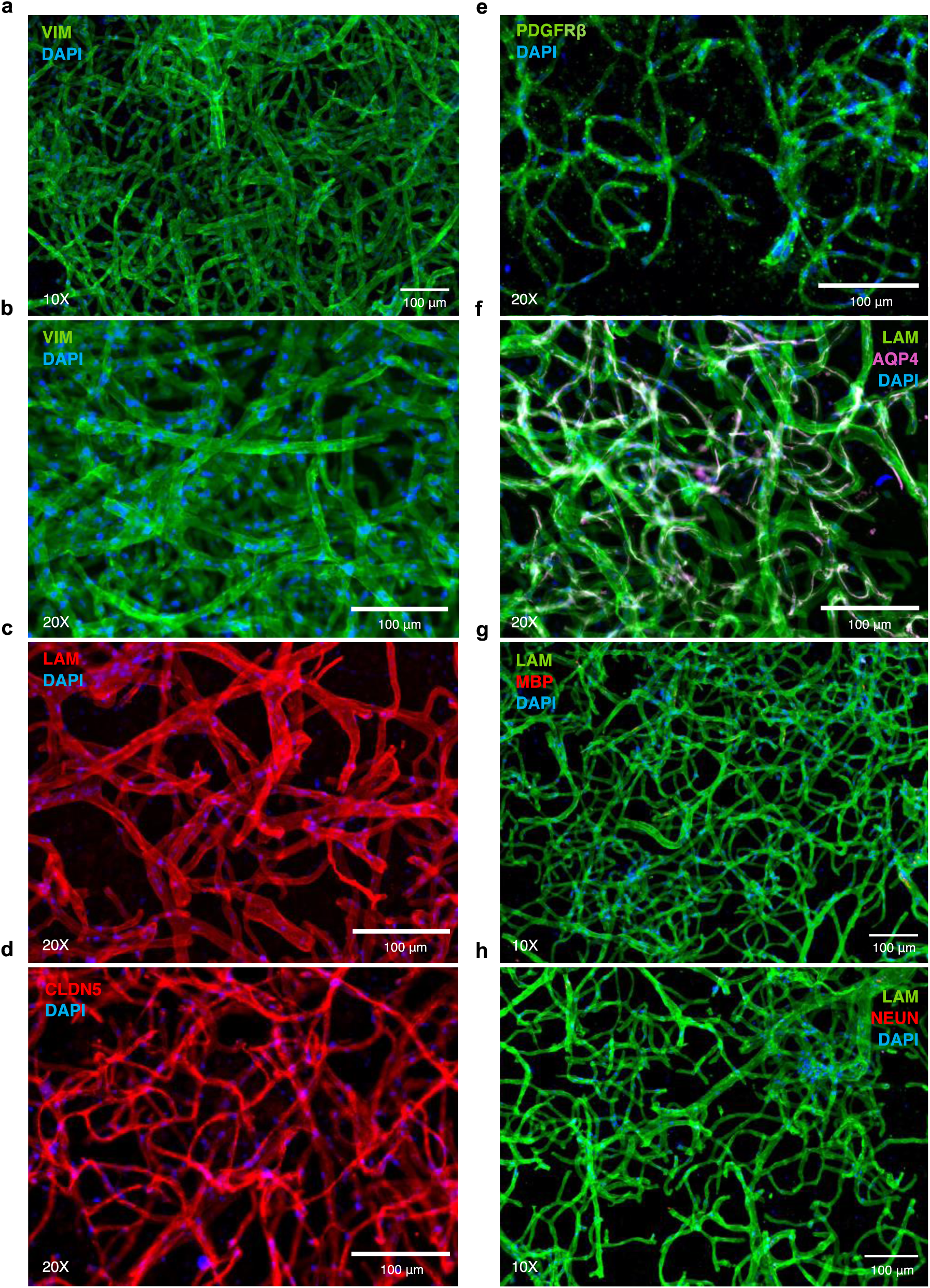
Isolated microvessels retain their *in situ* structure and express canonical markers. a-b) Endothelial and smooth muscle cells immunolabelled for vimentin (green). c) Extracellular matrix laminins expressed by vascular-related cell types are immunolabelled (red). d) Endothelial cell tight junctions immunolabelled for CLDN5 (red). e) Pericytes immunolabelled for PDGFRβ (green). f) AQP4 expressed by the vessel-facing astrocyte membrane is immunolabelled (magenta) and laminins (green). g) Oligodendrocyte marker MBP (red) shows an absence of immunoreactivity in microvessel preparations, similarly, h) neuronal marker NeuN (red) shows an absence of immunoreactivity in microvessel preparations. Nuclei are stained with DAPI (blue) in all micrographs.

Astrocytes serve multiple essential functions in supporting normal brain physiology^110^. This is, in part, due to the extension of astrocytic endfeet that surround approximately 99% of the cerebrovascular surface^111^ which, in conjunction with pericytes^112^, regulate expression of molecules that form the BBB including tight junction, enzymatic, and transporter proteins^113–115^. Although astrocytes were not co-isolated with microvessels, their perivascular endfeet are ensheathed within the same vascular basement membrane as endothelial cells and pericytes, making it possible that astrocytic endfeet remained attached after tissue homogenization. Because of this, we further observed astrocyte vascular coverage of isolated microvessels. AQP4, a water channel protein essential for the maintenance of osmotic composition and volume within the interstitial, glial and neuronal compartments^116, 117^, is expressed at the vessel-facing astrocytic membrane and superimposes the walls of isolated microvessels (Fig. 5f).

Finally, we examined possible contamination of our microvessel preparations by other cell types found within the brain. Immunolabelling for myelin basic protein (MBP; anti-Myelin Basic Protein antibody, BioLegend) was not detected (Fig. 5g); nor did immunolabelling for neuronal nuclear protein (NeuN; anti-NeuN antibody clone A60, Sigma-Aldrich) reveal the presence of neuronal elements (Fig. 5h). Thus, neurons and oligodendrocytes were consistently not observed to be co-isolated with microvessels. Together, these results indicate that the described protocol allows for the isolation and enrichment of structurally preserved brain microvessel fragments that are comprised of BMECs, astrocytic endfeet, pericytes, and tight junction proteins.

### 3.2. Computational deconvolution and characterization of transcriptomic data indicates high microvessel yield after isolation

To estimate the enrichment of our microvessel preparations, we performed computational deconvolution using calculated TPMs (Fig. 6a-b) using the BrainDeconvShiny tool (https://voineagulab.shinyapps.io/BrainDeconvShiny/). To demonstrate stability of outcome regardless of resources used, different iterations of deconvolution were performed utilizing single-nucleus datasets from Velmeshev et al. (2019; VL)^71^ and the MultiBrain (MB) composite signature from Sutton et al. (2022)^103^ as the reference signature, using the CIBERSORT v1.04 algorithm to deconvolute sample profiles and estimate cell type composition. Regardless of the approach used, majorly endothelial gene expression was returned, with an average of 94.92% (Fig. 6a) and 86.91% (Fig. 6b). Some contamination from neurons (1.29% and 4.66%, respectively) as well as negligible contamination from oligodendrocytes and microglia was observed. When using either dataset as the reference signature, limited presence of astrocytic genes was observed (1.16% and 6.11%, respectively), which may represent the contribution of astrocytic endfeet that cover the length of the neurovasculature. To estimate the multicellular composition of our microvessels at a finer resolution, computational deconvolution was performed a third time using the single-nucleus data generated by Yang et al. (2022)^65^, in which the different neurovascular cell type signatures were determined, as the reference signature (Fig. 6c). Averaged CPMs across nuclei of each cell type were used as input for the CIBERSORTX tool^105^, which estimated an average composition of 44.02% capillary and 37.62% SMC, along with much lower estimations for pericyte, arterial, venous, astrocyte and perivascular fibroblast genes (Fig. 6c). Although differentially distributed along the arteriovenous axis, both SMCs and pericytes are embedded within the vascular basement membrane^11^ and, therefore, it is unlikely that our isolated method preferentially selects one cell type over the other. Because of this, and the known similarity in molecular signature between SMCs and pericytes^118, 119^, as well as the high percentage of captured capillary segments (in which pericytes are predominantly observed)^120–123^, the surprisingly low estimation of pericyte genes may represent a limitation in comparing single-nuclei and bulk tissue datasets to one another. Critically, an average of 90.4% of the total TPMs across samples were assigned to NVU-constituent cell types.

**Figure 6.**
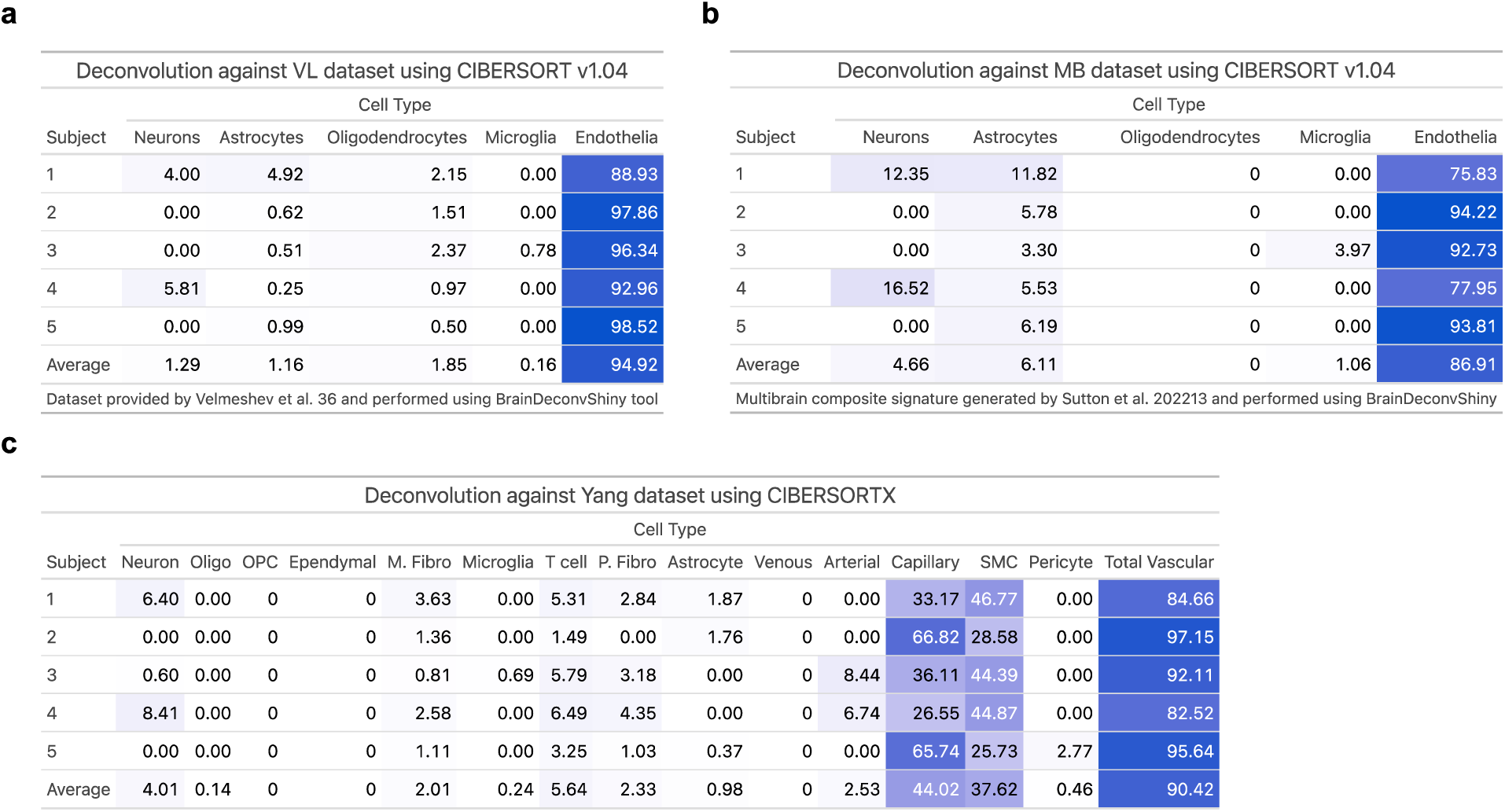
computational deconvolution indicates high microvessel yield after isolation. a-b) Using the BrainDeconvShiny tool, different iterations of computational deconvolution were performed to demonstrate stability of outcome when using our microvessel isolation preparations. a) The average expression from every control sample from the VL dataset was calculated and used as cell type signatures for deconvolution of our dataset using CIBERSORT v1.04. b) The MB (MultiBrain) is a composite signature generated by Sutton et al. (2022) by averaging the expression signatures of five datasets for five cell types (neurons, astrocytes, oligodendrocytes, microglia, and endothelia). MB was used as cell type signatures for deconvolution of our dataset using CIBERSORT v1.04. c) In-house analysis where the average expression from every control sample from the Yang dataset was calculated and used as cell type signatures for deconvolution of our dataset using CIBERSORTX. Heat tables generated using the gt package in R.

To explore our transcriptomic data, the top 10% of most highly expressed genes were designated (a total of 3968 genes) and used as input for over-representation analysis (ORA) using the enrichR package in R, selecting the “Descartes Cell Types and Tissue 2021” database to identify gene sets that are statistically over-represented (Fig. 7a-b and Supplementary Tables 4). The threshold value of enrichment was selected by a p-value <0.05 and, as shown, over-represented genes were dramatically enriched for vascular-related terms, such as “Vascular endothelial cells in Cerebellum” and “Vascular endothelial cells in Cerebrum”. Genes behind enriched terms were extracted and the expression of a subset of known brain endothelial, pericyte, astrocytic and smooth muscle genes in isolated microvessels were examined (Fig. 7c-i). As expected, microvessels had increased expression of canonical endothelial genes such as CLDN5, CDH5, SLC2A1, ABCB1, VWF, and MFSD2A; with highest expression in endothelial genes ACTG1, B2M, BSG, EEF1A1, HLA-B, HLA-E, SPARCL1, TMSB10, and VIM (Fig. 7a).

**Figure 7.**
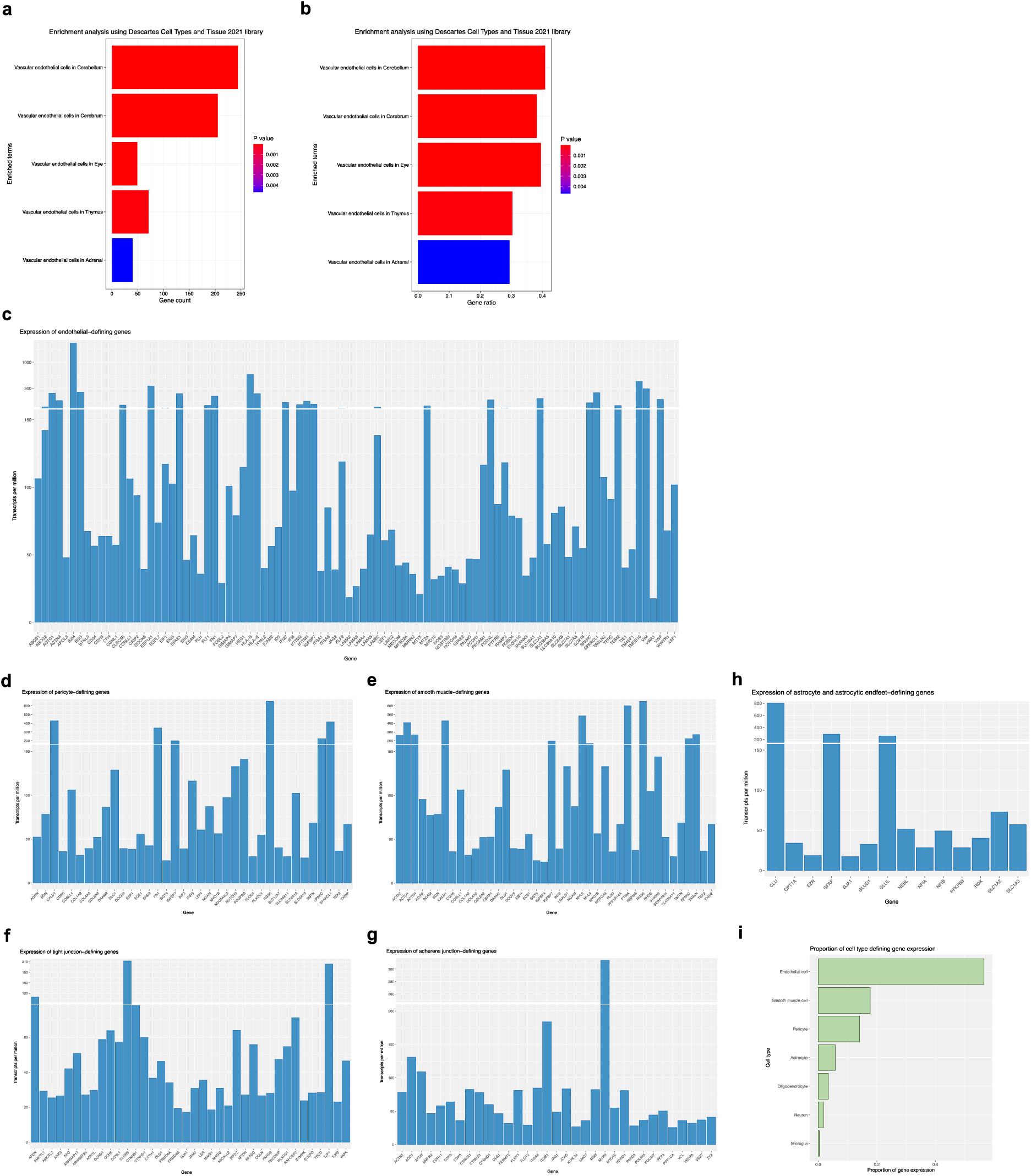
Analysis of microvessels gene expression indicates overrepresentation of neurovascular-related gene expression. a-b) Enrichment analysis of top 10% most highly expressed genes (3968 genes) returns predominantly vascular-related terms. Over-representation analysis (ORA) using the enrichR package in R and the “Descartes Cell Types and Tissue 2021” database was used to identify gene sets that are statistically over-represented. The threshold value of enrichment was selected by a p-value <0.05, indicating over-represented genes were significantly enriched for vascular-related terms. a) Count for genes in our dataset that are present in returned gene sets. b) Ratio for genes in our dataset that are present in returned gene sets, determined by the total number of genes in each set. c-i) Isolated microvessels have increased expression of canonical neurovascular-related genes. c) Bar plot showing TPMs for endothelial-defining genes. Highest expression found in endothelial genes B2M, BSG, FLT1, IFITM3, MT2A, SLC2A1, VIM, VWF. d) Bar plot showing TPMs for pericyte-defining genes. Highest expression was detected in pericyte genes CALD1, FN1, IGFBP7, RGS5, SPARCL1. e) Bar plot showing TPMs for smooth muscle cell-defining genes. Highest expression was detected in smooth muscle genes ACTG1, ACTN4, MYL6, PTMA and TAGLN. f) Bar plot showing TPMs for tight junction-defining genes and g) Bar plot showing TPMs for adherens junction-defining genes. Several genes encoding for junctional proteins are found in the top 10% of most highly expressed genes, including CLDN5, CTNNB1, CTNND1, OCLN, JAM1, TJP1, and TJP2. h) Bar plot showing TPMs for astrocyte and astrocytic endfeet-defining genes. Expression of astrocytic genes were predominantly limited to markers of astrocytic processes or endfeet, namely CLU, GFAP, and GLUL. i) TPMs from neurovascular-related genes were summarized according to cell type expression, demonstrating overrepresentation of endothelial, smooth muscle cell, and pericyte genes. Bar plots generated using the ggplot package in R.

Additionally, there was enrichment for other neurovascular cell types, as suggested by canonical pericyte genes PDGFRβ, MCAM, RGS5, AGRN, and NOTCH3 (Fig. 7b); as well as canonical smooth muscle genes ACTA2, MYL6, MYL9, TAGLN, and LGALS1 (Fig. 7c). Highest expression was detected in pericyte genes CALD1, FN1, IFGBP7, RGS5, SPARC and SPARCL1 (Fig. 7b); as well as smooth muscle genes ACTG1, ACTN4, CALD1, MYL6, and PTMA (Fig. 7c). Complimentary to immunophenotypic characterization of isolated microvessels, several genes whose products are involved in junctional complex maintenance and organization (tight junctions, Fig. 7f; adherens junctions, Fig. 7g) are found in the top 10% of most highly expressed genes, for e.g., CLDN5, CTNNB1, CTNND1, ITGB1, JAM1, OCLN, TJP1, and TJP2; with additional vascular makers expressed at lower transcripts per million are present throughout the entire dataset. Expression of astrocytic genes were observed to a lesser extent, a number of which have been previously validated as markers of astrocytic processes or endfeet^124–126^, namely EZR, GJA1, RDX, SLC1A2, and SLC1A3 (Fig. 7h). To better visualize the proportion of expression contributed by these genes, TPMs were summarized by cell type over the total number of TPMs (Fig. 7i), demonstrating overrepresentation of endothelial, smooth muscle cell, and pericyte genes in samples.

Intriguingly, results from ORA reveal genes that are of interest in numerous disease contexts. Enriched in the described dataset is gene CLDN5, an indispensable junctional protein for the correct organization of tight junctions and maintenance of BMEC integrity^108^, which was previously reported to be downregulated in the nucleus accumbens of depressed suicides^127^ and targeted altered epigenetic regulation via histone deacetylase 1 (HDAC1)^128^. Enriched pericyte genes TXNIP, RUNX1T1, ITGA1 and DOCK9 are also reported to be differentially expressed in Schizophrenia^129^. Similarly enriched in this dataset are angiogenic growth factors EGFL7, FLT1, VWF and antigen-presentation machinery B2M and HLA-E, all of which are upregulated in a subpopulation of angiogenic BMECs that are induced in subjects with Alzheimer’s disease^130^, suggesting a compensatory angiogenic and immune response in AD pathogenesis. Likewise, endothelial genes PICALM, INPP5D, ADAMTS1, PLCG2 that are found in the top 10% of the microvessel dataset are differentially expressed in Alzheimer’s disease^65^. Recent breakthroughs in deciphering the underlying etiology of Huntington’s disease reveals aberrant downregulation of endothelial ABCB1, ABCG2, SLC2A1 and MFSD2A as well as mural (which constitutes pericytes and vascular smooth muscle cells) PDGFRB, SLC20A2 and FTH1^64^ *–* mutations in which are known to cause HD-like syndromes with primary pathology localized in the basal ganglia^131, 132^. Indeed, brain microvessel datasets such as these integrated with others, like as those from experimental models, will expedite our understanding of neurovascular contributions to mood disorders and neurodegenerative disease, and even further propagate hypothesis generation for established vascular diseases, such as white matter vascular dementia in which each cell type of the NVU exhibits a specific disease-associated expression signature^133^.

### 3.3. High correspondence between generated transcriptomic data and published neurovascular dataset

As a means to assess our isolation method, our top 10% most highly expressed genes were juxtaposed to validated neurovascular cell type-defining markers, as designated by Garcia et al. (2022)^64^ based on sequencing of 4,992 and 11,689 vascular nuclei from *ex vivo* and postmortem brain tissue, respectively. Results indicated substantial overlap between the three datasets for all NVU-constituent cell types (Fig. 8), including arteriole-defining genes (Fig. 8a), capillary-defining genes (Fig. 8b), venule-defining genes (Fig. 8c), arteriolar SMC-defining genes (Fig. 8d), pericyte-defining genes (Fig. 8e), venular SMC-defining genes (Fig. 8f), and perivascular fibroblast-defining genes (Fig. 8g). Interestingly, overlap was consistently greater between the two postmortem datasets.

**Figure 8.**
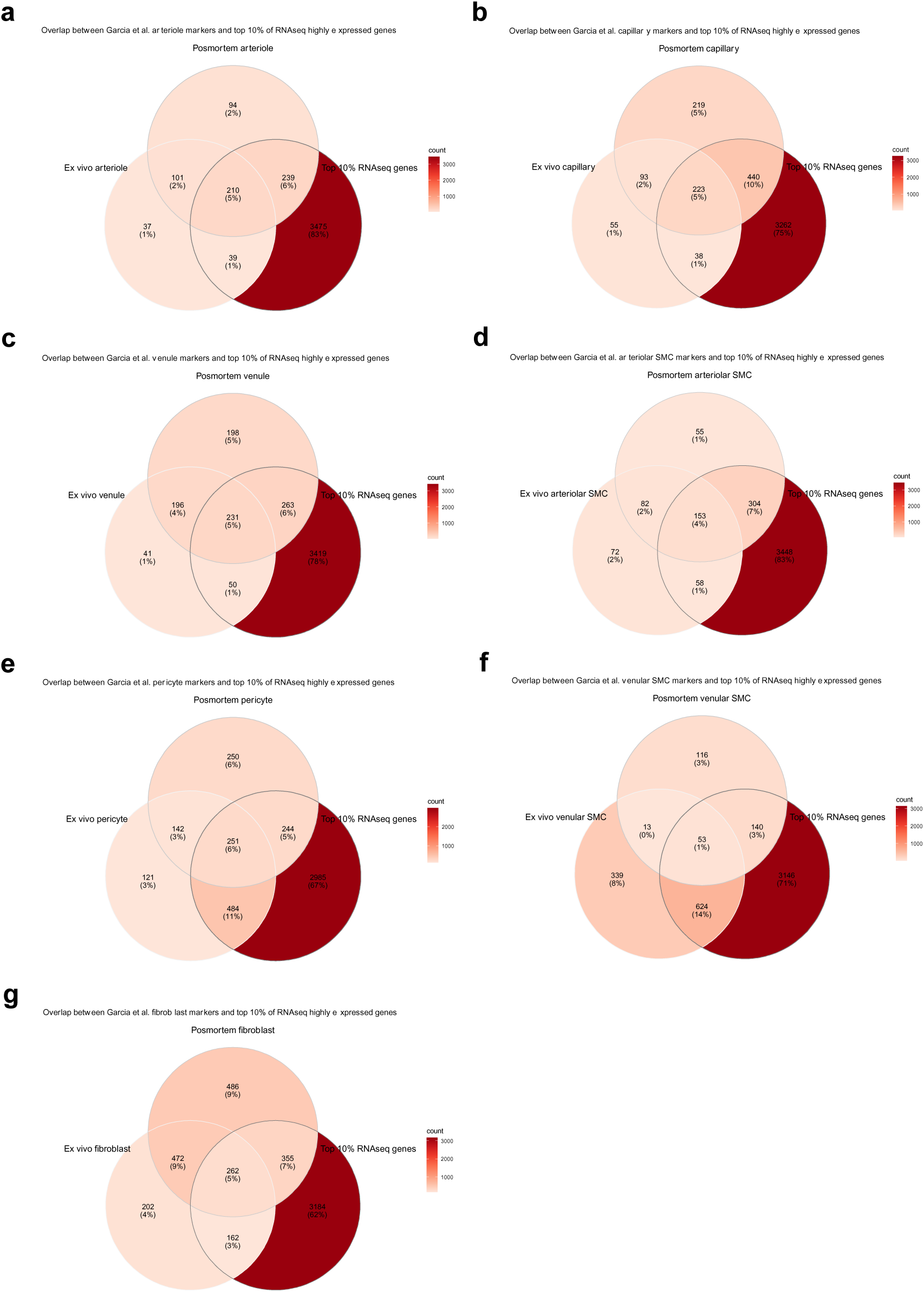
Comparison of RNA sequencing data to published neurovascular dataset. a-g) Overlap between top 10% of highly expressed genes from our RNA sequencing data and published NVU dataset. Validated neurovascular cell type markers were obtained from Garcia et al. (2022) postmortem and human *ex vivo* single-nuclei sequencing datasets (found under supplementary table 2) and compared for potential overlap with the top 10% of most highly expressed genes from our data. Results indicated significant overlap between the three datasets for all NVU-related cell types. a) Arteriole-defining genes. b) Capillary-defining genes. c) Venule-defining genes. d) Arteriolar SMC-defining genes. e) Pericyte-defining genes. f) Venular SMC-defining genes. g) Fibroblast-defining genes. Venn diagrams generated using the ggVennDiagram package in R.

### 3.4. Proteomic characterization of isolated brain microvessels

While transcript level may show positive correlations with protein level, protein abundance should not necessarily be inferred from RNA sequencing counts^134, 135^, hence, interrogation of the highly dynamic proteome is needed to speculate the functional consequences of changes in protein expression. We sought to interrogate NVU-specific proteomic signatures using total protein extracted from isolated microvessels. Microvessel-enriched pellets were prepared from 3 frozen vmPFC grey matter samples microdissected from healthy individuals who died of peripheral diseases or natural events (Table 1). Although resuspended and strained pellets yield sufficient protein needed to perform LC-MS/MS (validated, data not shown), straining was omitted in favour of protein extraction directly from pellets to maximize microvessel material input for proteomic interrogation. Using Tandem Mass Tag (TMT) isobaric labeling and sample fractionation of peptides, global, relative quantitation of a total of 1638 individual proteins were detected from microvessel-enriched pellets (quality parametres shown in supplementary Fig. 2). Importantly, there was significant overlap between transcriptomic and proteomic output, with 1635/1638 (99.8%) of detected proteins likewise identified in the transcriptomics data (albeit no corresponding transcripts were detected for proteins F9, DCD, and SERPINB12); and with 1034/1638 (62.5%) proteins found in the top 10% of most highly expressed transcripts, resulting in an overall 23% overlap between high expressors in both datasets (Fig. 9a).

**Figure 9.**
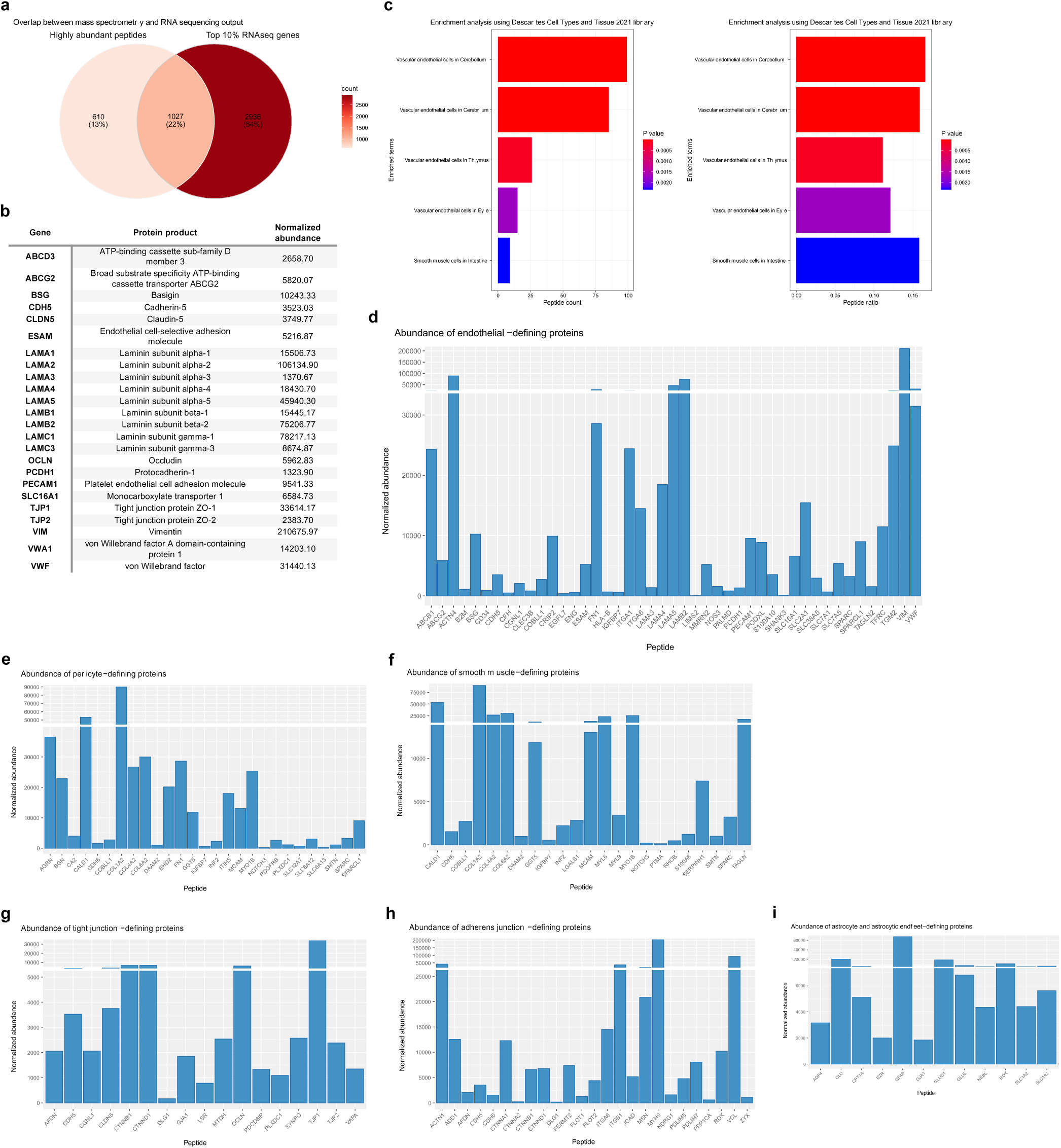
Overlap between transcriptomic and proteomic data output. a-i) There is substantial overlap between transcriptomic and proteomic data output. a) 1635/1637 (99.9%) of proteins likewise identified in the transcriptomics data and with 1024/1637 (62.6%) proteins found in the top 10% of most highly expressed genes. b) Several canonical BMEC markers were detected with high normalized average abundance. c) Over-representation analysis using the enrichR package in R and the “Descartes Cell Types and Tissue 2021” database was used to identify sets that are statistically over-represented. Genes corresponding to proteins detected during LC-MS/MS were used as input and the threshold value of enrichment was selected by a p-value <0.05. Output indicated over-represented proteins were significantly enriched for vascular-related terms. a) Count for peptides in our dataset that are present in returned sets. b) Ratio for peptides in our dataset that are present in returned sets, determined by the total number in each set. d) Bar plot showing normalized abundance for endothelial cell-defining proteins. e) Bar plot showing normalized abundance for pericyte-defining proteins. f) Bar plot showing normalized abundance for smooth muscle cell defining-proteins. g) Bar plot showing normalized abundance for tight junction defining-proteins. h) Bar plot showing normalized abundance for adherens junction-defining proteins. i) Bar plot showing normalized abundance for astrocyte and astrocytic endfeet-defining proteins. Bar plots generated using the ggplot package and Venn diagram using the ggVennDiagram package in R.

Proteins known to be expressed by vascular-associated cell types and perivascular extracellular matrix were positively identified in all 3 samples. Several canonical endothelial markers were detected with high peptide abundance, including vimentin, several protein subunits of laminin (LAMA2, LAMC1, LAMB2, LAMA5, LAMA4, LAMA1, LAMB1, LAMC3, LAMA3), OCLN, TJP1, TJP2, VWF, VWA1, BSG, PECAM1, Monocarboxylate transporter 1 (SLC16A1), broad substrate specificity ATP-binding cassette transporter (ABCG2), ESAM, CLDN5, CDH5, ATP-binding cassette sub-family D member 3 (ABCD3), and Protocadherin-1 (PCDH1) (normalized average abundance for these proteins are listed in Fig. 9b). As expected, enrichment terms returned by ORA (using corresponding gene names as input) were predominantly vascular-associated (Fig. 9c and supplementary Table 5) and normalized abundance of known brain endothelial, pericyte, astrocytic and smooth muscle cell proteins were high (Fig. 9d-i).

The BBB poses as a major pharmacological barrier as BMECs express a vast array of enzymes and transport systems that facilitate brain uptake processes of essential nutrients and neuroactive agents across the BBB^136^, and therefore control the rate and extent to which drugs are able to reach the brain parenchyma via the transcellular pathway^137^. There is a pressing need for greater knowledge surrounding the expression and functionality of these systems at the human BBB as the majority of data comes from either *in vitro* cell culture or animal studies, making *in vitro* to *in vivo* or interspecies scaling less reliable. Analyses revealed high abundance of SLC2A1/GLUT1, a transmembrane protein responsible for the facilitated diffusion of glucose^138^, and the two glutamate transporters SLC1A2/EAAT2 and SLC1A3/EAAT1 in brain microvessels. Transporters SLC7A5/LAT1 and SLC3A2/4F2hc, which supply the brain with large neutral amino acids^139–141^, and SLC16A1/MCT1 and SLC16A2/MCT8, which are involved in the transport of monocarboxylates^142^ and T3 thyroid hormone^143^ at the BBB, respectively, are also found in the described proteomic dataset. Also found are ABCB1/P-glycoprotein and breast cancer-related protein ABCG2/BCRP, the principal ABC efflux transporters^144–146^ that limit entry of drug candidates, toxic compounds, as well as xenobiotics from the central nervous system^147, 148, 149, 150, 151, 152^. Due to major species-specific differences in transporter expression profiles, there is utility in obtaining absolute protein amounts as they may elucidate the contribution of each transporter in facilitating the entry of endogenous substances and nutrients like glucose, glutamate, and amino and fatty acids into the brain, in addition to drugs and other xenobiotics that exploit these mechanisms^153^.

### 3.5. Further deconstruction of isolated brain microvessels using FANS

The use of frozen brain tissue demands sorting of target nuclei as opposed to intact cells and, therefore, requires the use of nuclear fluorescent tags to facilitate isolation of endothelial nuclei. Expression of transcription factor ETS-related gene (ERG) has been previously validated as an endothelial-specific nuclear marker^98, 154–156^. As an additional testament to the versatility in downstream usability of the described method, resuspended microvessels subjected to a 2-hour incubation with anti-ERG antibody under rotation is sufficient for vascular disassembly and to expose the ERG epitope for endothelial nuclei immunolabelling (Fig. 10a-b). Importantly, enzymatic digestion is not needed as previously described^78, 157, 158^ and, in combination with the appropriate gating strategy adjusted to collect endothelial nuclei, is sufficient for the purification of endothelial nuclei from frozen postmortem brain tissue (data not shown). The potential to isolate ERG^+^ endothelial nuclei from postmortem microvessels under different physiological conditions^159–165^ is an advantage of our approach and addresses a growing interest in the use of primary BMECs, circumventing phenotypical or gene expression changes induced in primary BMEC cultures by by prolonged adherence steps used in other isolation protocols^166, 167, 168, 169^.

**Figure 10.**
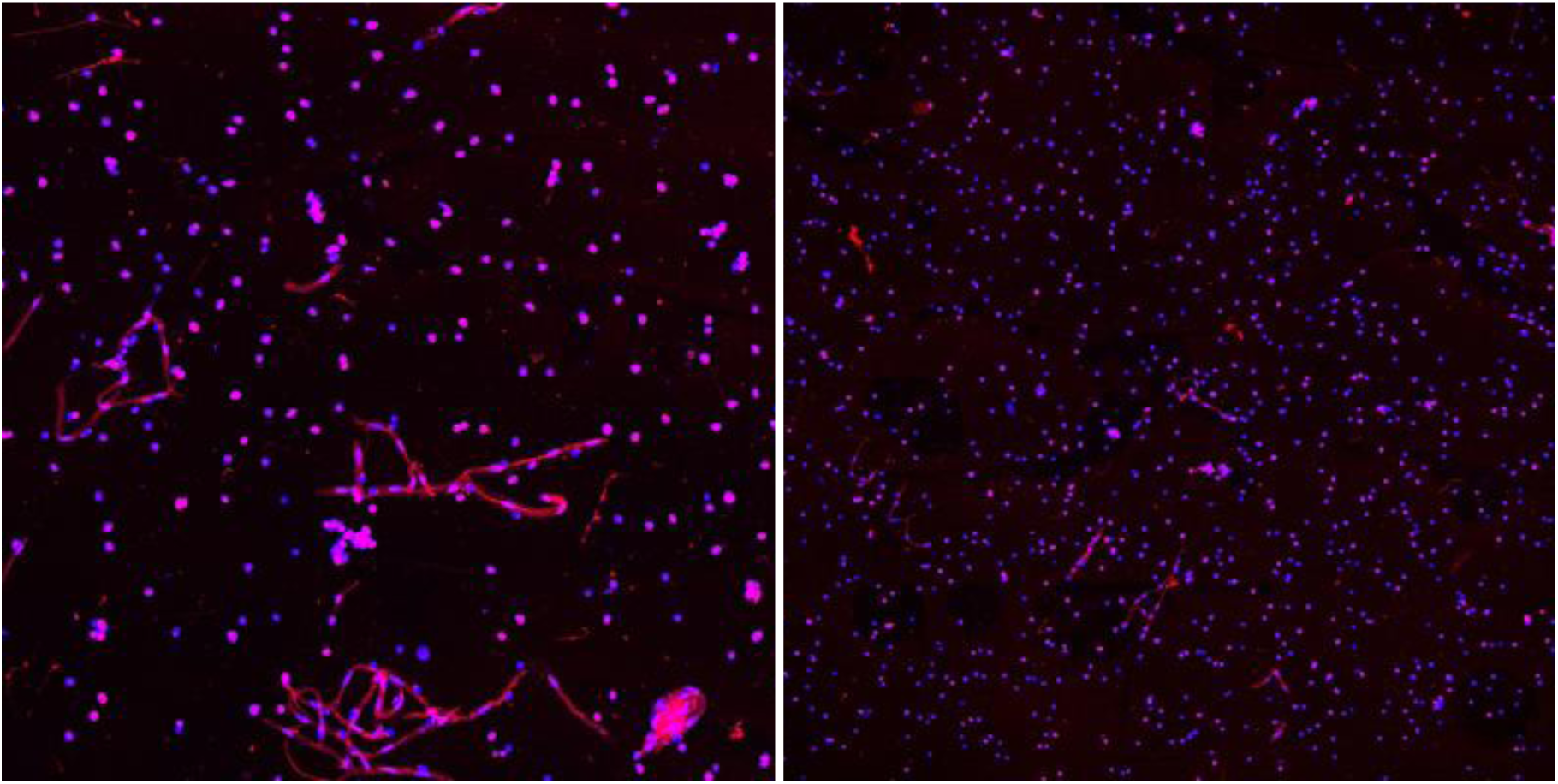
Isolated microvesselscan be used to further sort endothelial nuclei. a-b) Isolated microvessels can be used to further sort endothelial nuclei. a-b) Dissociated endothelial nuclei immunolabelled with ERG conjugated to Alexa-647 antibody prior to FANS protocol. Resuspended microvessels subjected to a 2-hour incubation with anti-ERG antibody under rotation is sufficient for immunolabelling of endothelial nuclei prior to FANS sorting.

## 4. Discussion

We describe a singular, standardized protocol to enrich and isolate microvessels from archived snap-frozen human with the ability to apply the same protocol to frozen mouse cerebral cortex. Integral to this method are three factors that confer gentility and simplicity: 1) the correct molarity of the sucrose-based buffer to separate and cushion microvessels during centrifugation-separation, 2) centrifugation-separation of microvessels from the rest of tissue homogenate occurs in a 10 ml volume, which facilitates the formation of a microvessel-enriched pellet as heavier structures reach the bottom of the 15 ml falcon tube, and 3) the limited number of wash steps that minimize eventual damage done to the integrity of microvessel fragments. Microvessel enrichment and stability was assessed by chromogenic staining of microvessels using BCIP/NBT substrate and, through immunophenotypic characterization, we show that isolated microvessel fragments are comprised of NVU cellular components including BMECs, astrocytic endfeet, pericytes, as well as tight junction protein complexes. The demonstrated gentility and simplicity of the approach, in turn, confer versatility in the high-throughput techniques that can be successfully utilized downstream to microvessel isolation, as demonstrated here with RNA sequencing and LC-MS/MS.

This protocol should have excellent reproducibility in isolating intact microvessels from any region of the adult human brain. It should be noted, however, that the current version of the protocol may need some adjustments if myelin content of brain samples used is high. As reported, microvessels isolated using the described method are in high yield in the absence of other contaminating cell types. However, limited expression of neuronal markers is observed during computational deconvolution of RNA sequencing data. Whether a minority of neurons are co-enriched with microvessels remains unclear, as co-enrichment of neurons was not observed during immunophenotypic characterization of isolated microvessels. While some co-enrichment is consistent with the technically unavoidable capture of some parenchymal cells when utilizing a method suitable for frozen postmortem brain tissue (which necessitates as few and as gentle steps as possible), the human BBB atlases generated by Yang et al. (2022) and Garcia et al., (2022) reveal that “canonically neuronal” genes are also expressed in vascular-associated cell types. Therefore, it is possible that counts for such genes indeed originate from vascular-associated cells and not neurons. Another limitation is that the protocol enriches for but does not purify microvessels, so a limited proportion of contamination from non-neurovascular cell types might be found. Additionally, representation of all vascular-associated cell types might not be uniform, as different cell types may vary across brain regions and even brain subregions, and different cell types may be differentially susceptible to steps taken during RNA or protein extraction. However, it is reasonable to assume that the proportions of vascular-associated cell types found within isolated microvessels represent the natural multi-cellular composition of vasculature within the brain. Inevitably, studies that make use of isolated human microvessels will encounter high inter-subject variation in both RNA and protein yield as well as experimental read-out. Large cohorts with matched subjects (if studying disease and/or sex differences) would be required to power a study investigating how brain vascular gene or proteomic expression varies with factors such as brain region, age, or disease.

Although sensitivity of mass spectrometers has been greatly improved over the years^170–174^, as well as peptide separation for untargeted proteome analysis^172, 173, 175^, the literature consistently shows that high output from mass spectrometry relies on a fine balance between sample quantity as well as sample complexity. Because protein extracted from a structural unit, such as microvessels, as opposed to bulk tissue could be considered a sample lacking complexity, it is reasonable to assume that this relays to lower output, which may be further impacted by several low signal-to-noise events that result in unidentified peptides^176^ and a more limited dynamic range of peptide detection^174^. An example of this are low-molecular weight cytokines which are released in diluted concentrations by virtue of their high biological activity^177^.

Our successfully generated datasets promise that future datasets generated from microvessels isolated by the described method, when compared to those isolated from mice, have the potential to improve our understanding of species-specific differences in brain vascular gene expression, which is key to assess potential limitations of mouse models for brain vascular and BBB development and disease. Species differences in disease-affected neuronal and myeloid gene expression have been characterized by single-cell or single-nucleus sequencing^178–180^, but such differences in neurovascular gene expression have only just begun to be comprehensively analyzed, and solely in Alzheimer’s disease and Huntington’s disease^64, 65^. Despite limited data, it has become evident there are striking differences, with one study reporting 142 mouse-enriched genes, including Vtn, Slco1c1, Slc6a20a, Atp13a5, Slc22a8, and 211 human-enriched genes, including SLCO2A1, GIMAP7, and A2M^181^. Yang et al. (2022)^65^ further reported hundreds of species-enriched genes in BMECs and pericytes^65^, finding that BECs and pericytes exhibit the greatest transcriptional divergence in several vascular solute transporters (for e.g., GABA transporter SLC6A12) and genes of disease and pharmacological importance^65^. This observation was corroborated by Garcia et al. (2022) ^64^, who further detailed that species-specific DEGs were strongly enriched for marker genes of vascular-associated cell types, indicating that cell type identity markers were among those that vary the most between species^64^.

In sum, the isolated brain microvessel is a robust model for the NVU and can be used to generate a variety of highly dimensional datasets. The availability of characterized human neurovascular transcriptomes and proteomes can aid investigators in identifying potential roles for BMECs and pericytes in the pathogenesis of neurological and psychiatric conditions, and should allow others to quickly assess the expression of molecules with potential relevance to drug delivery (e.g., SLCs, ABCs, and large molecule receptors) and disease in the human brain vasculature.

## Acknowledgements

The authors have no competing interests to declare. This study was funded by an ERA NET Neuron grant and a CIHR Project grant to N.M. M.W. and R.R. respectively received scholarship and fellowship support from the FRQ-S. The Douglas-Bell Canada Brain Bank is supported in part by platform support grants from the Réseau Québécois sur le Suicide, les Troubles de l’Humeur et les Troubles Associés (FRQ-S), Healthy Brains for Healthy Lives (CFREF), and Brain Canada. The present study used the services of the Molecular and Cellular Microscopy Platform (MCMP) at the Douglas Research Centre. The authors would like to thank the expert help of Douglas-Bell Canada Brain Bank staff members (J. Prud’homme, M. Bouchouka, D. Mirault, V. Lariviere, A. Baccichet), the technology development team at the McGill University and Genome Quebec Innovation Centre, and the Segal Cancer Proteomics Centre at the Lady Davis Institute. The authors would also like to thank Dr. Ghazal Fakhfouri for rearing and providing the mice used in this study.

