## supplementary Table for "Universal method for the gentle isolation of intact microvessels from frozen tissue: a multiomic investigation into the neurovasculature"

**Supplementary data**

**Supplementary Table 1: Efficacy of DEPC as a cost-effective RNAse inhibitor during microvessel isolation**

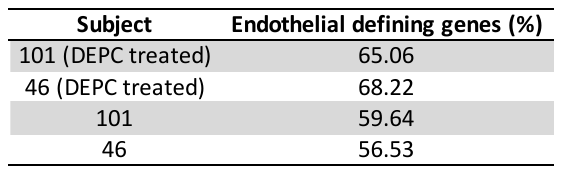

If transcriptomic techniques are to be used downstream to microvessel isolation, it is recommended to use DEPC-treated water as opposed to Millipore water, as DEPC acts as a RNAse inhibitor. To demonstrate the utility of DEPC as a cost-effective RNAse inhibitor when homogenizing brain tissue in large volumes of buffer, the use of DEPC-treated water to prepare the Homogenization Buffer was compared directly against the same experimental workflow without DEPC treatment. RNA was extracted from isolated microvessels using the Single Cell RNA Purification Kit (Norgen Biotek Corp., Thorold, Ontario) and sequencing libraries were built using the SMARTer Stranded Total RNA-Seq Kit v3 - Pico Input Mammalian (Takara Bio Inc., Shiga, Japan). Libraries were sequenced on the MiSeq platform (Illumina, Inc.; California, United States) and raw sequencing data processed using inhouse pipeline. Computational deconvolution was performed using the DeconRNAseq R package and the previously published dataset provided by Nagy & Maitra et al. (2020, **32341540). As shown here,** DEPC treatment preserves an approximate 10% of transcripts that are otherwise lost when processing samples for RNA-sequencing.

**Supplementary Table 2: Measurements of collected microvessel-enriched pellets**

| **Subject** | **Weight of tissue microdissection (mg)** | **Macroscopic observations** | **Weight of wet microvessel-enriched pellet (mg)** | **RNA conc. (ng/ul)** | **RIN of microvessel-enriched pellet** |
| --- | --- | --- | --- | --- | --- |
| **1** | 106.8 | Normal appearance | 94.55 | 11.3 | 4.3 |
| **2** | 101.28 | Normal appearance | 84.07 | 9.58 | 2.9 |
| **3** | 103.07 | Normal appearance | 77.02 | 6.75 | 5.2 |
| **4** | 104.26 | Normal appearance | 77.53 | 4.65 | 4.3 |
| **5** | 103.88 | Normal appearance | 72.72 | 10.4 | 4.4 |
| **Average** | 103.86 |  | 81.18 | 8.54 | 4.22 |

Information regarding microvessel samples subjected to RNA sequencing, including weight of tissue microdissection, weight of collected microvessel-enriched pellet, concentration of RNA extracted from pellets, as well as RIN of extracted RNA.

**Supplementary Table 3: Anticipated methodological issue and potential solutions**

| **Issue** | **Possible reason** | **Solution** |
| --- | --- | --- |
| Small clumps of tissue are visible in lysate after homogenization | Too much tissue is homogenized at once | This protocol has been optimized to isolate microvessels specifically from 100 mg of frozen brain tissue, however up to 120 mg of tissue at once has been used with success. It is best to keep tissue samples within this range. Note that tissue samples microdissected into several small pieces might equilibrate/homogenize more readily. |
|  | Wrong homogenization program is used | Due to variation in brain tissue quality and integrity at collection, PMI, and storage practices across brain banks, it is possible that another homogenization program is better suited. However, the different programs available using the gentleMACS™ Dissociator have been exhaustively explored during optimization of this protocol, with the most appropriate deemed to be Lung 02.01 |
| Pellet is smaller than expected or not present | Poor tissue quality | When possible, tissue samples with high RIN (>5) should be used |
|  | Tissue not given time to equilibrate in buffer before homogenization | Gently agitating tissue in buffer for 30 seconds before homogenization will encourage thawing and osmotic equilibrium that will allow for uniform dissociation of tissue. |
|  | Loss of vessel fragments to the myelin layer | This protocol has been optimized for predominantly grey matter tissue and high white matter content should be avoided. While samples with some white matter content have been regularly used with success, higher levels of myelin may affect pellet yield. This is because dissociated myelin will clump together and rise to the top of the lysate during centrifugation, potentially trapping microvessels that would otherwise travel to the bottom of the tube |
| Pellet is larger than expected | Myelin layer not well separated from the microvessel pellet/contamination of pellet with fat | Use of a 15 ml falcon tube during 30-minute centrifugation allows sufficient distance and time for the correct structures to travel downwards (microvessels) and correct structures to travel upwards (myelin). As mentioned above, samples with lower myelin content should be used |
|  | Tissue insufficiently homogenized | See suggestions listed for first problem |
| Resuspended pellet does not filter through the cellular strainer | Mesh is clogged | Because of the low pore size, the resuspended pellet may not easily pass through the strainer. Thus, it is recommended to pipette sample slowly and gradually through the 35μm strainer |
|  |  | Use vacuum-aspiration underneath the strainer cap to encourage filtration. Vacuum-aspiration should be applied lightly and in pulses |

**Supplementary Table 4: Returned enrichment terms (analysis of RNA sequencing data)**

| **Term** | **Overlap** | **P-value** | **Adjusted P-value** | **Odds Ratio** | **Combined Score** | **Genes** |
| --- | --- | --- | --- | --- | --- | --- |
| **Vascular endothelial cells in Cerebellum** | 244/597 | 3.72303406628903E-33 | 5.91962416539956E-31 | 2.91504844200252 | 217.668909197257 | IFITM3;IFITM1;IFITM2;SPARC;TFRC;ICAM2;MT1X;IFIT1;IFIT3;CAPNS1;SOX17;SOX18;GJA4;BSG;SLC12A7;MFSD2A;PRKCH;SLC6A13;SLC6A12;ACTN4;HSPG2;TM4SF1;SIGIRR;SLC5A6;CLDN5;BCAM;IFI27;DAAM2;TAGLN2;MYL9;MT1E;ABCB1;CFH;EPAS1;USHBP1;PDGFB;IQGAP1;GATA2;APCDD1;KIAA0040;MYL12A;MYL12B;PLAC9;RRAS;S100A16;SERPINH1;S100A10;WWTR1;FOXF2;GADD45A;MCAM;FN1;IGF2;GNG11;HSPA12B;FOSL2;EHD2;OCLN;SMTN;ID1;NOSTRIN;ID3;ITM2A;CD320;ACVRL1;CLIC5;COL18A1;ROBO4;SYNM;TINAGL1;TGFB1I1;HBB;PTH1R;LRRC32;PEAR1;SHE;C1QTNF1;CSRP1;MECOM;HYAL2;TNKS1BP1;TIMP1;CD34;LMCD1;CGNL1;ZNF366;MAP4K2;EDN1;ANXA1;EDN3;HEG1;TSC22D3;BGN;SLC39A10;HBA2;RHOC;DUSP6;RHOB;SLC7A5;INF2;VWA1;GPRC5C;VAMP5;ENG;VAMP3;FOXC1;RAMP2;TAGLN;AHNAK;ADCY4;NDUFA4L2;IFI6;CRIP2;FSTL1;SLC7A1;CRIP1;AKAP2;FAM129B;TCEAL8;FENDRR;FGD5;FOXS1;SAMM50;APOL3;SLC16A1;PLXDC1;FKBP1A;DLC1;CCDC3;AGRN;PLOD1;GIMAP6;ETS1;GIMAP7;CTGF;CDH6;FOXQ1;CDH5;RGS5;EFEMP2;RGS3;PLS3;SLC39A8;MAP3K6;TGM2;PDGFRB;IGFBP4;LMO2;EMCN;COL4A2;COL4A1;SLC7A5P2;ERG;FKBP9;PHLDB2;KANK3;ABCG2;NOTCH3;SHC1;NOTCH4;ECSCR;ITPR3;SLC38A11;COX7A1;HIGD1B;NPAS2;SLC9A3R2;MUSTN1;CALD1;PODXL;COBLL1;IGFBP7;SLC38A3;SLC38A2;SLC38A5;BTNL9;GGT5;RBPMS;CAV2;CAV1;ARHGAP29;ISG15;PTPRB;RGCC;MOB2;SLCO1A2;BHLHE40;ESAM;ST6GALNAC1;FLT1;SLC2A1;ECE1;NRARP;LAMC1;CYR61;GPER1;GRASP;EGFL7;KLF13;VWF;ITGA1;ARAP3;LSR;SHROOM1;ACTA2;FGR;ADORA2A;COL6A2;MMRN2;SHANK3;ITIH5;SLFN5;LEF1;SEMA3G;ATP10A;PRELP;SLC3A2;UACA;TTN;DLL4;CLEC3B;PALMD;CCDC85B;TMEM204;HES1;GBP1;GBP4;PPP1R14A;MYO10;LAMB2;NOS3;TIE1;KLF4;KLF2;TBX18;TBX2;MYO1B;PINK1;MYO1C;ABHD17A;RAB13;NPIPB4;GPD1L;TEK |
| **Vascular endothelial cells in Cerebrum** | 205/536 | 1.19397686280788E-23 | 9.49211605932263E-22 | 2.58841962926221 | 136.622398437087 | IFITM3;IFITM1;SPARC;TFRC;MT1X;IFIT3;PREX2;MT2A;LGALS1;SOX17;SOX18;SLC12A7;PRKCH;SLC6A13;SLC6A12;ACTN4;TM4SF1;SIGIRR;SLC5A6;BCAM;RRAGA;IFI27;RBP1;SERPING1;MT1E;ABCB1;CFH;EPAS1;USHBP1;GATA2;PLAC9;RRAS;S100A16;SERPINH1;S100A10;JAG2;WWTR1;PRELID1;FOXF2;MCAM;FN1;IGF2;HSPA12B;FOSL2;EHD2;OCLN;RPS28;COL1A2;ID1;NOSTRIN;ID3;ITM2A;CD320;ACVRL1;CLIC5;COL18A1;ROBO4;SYNM;TINAGL1;HBB;PTH1R;LRRC32;C1QTNF1;MECOM;HYAL2;TNKS1BP1;CD34;LMCD1;CGNL1;ZNF366;EDN1;ANXA1;EDN3;HEG1;BGN;SLC39A10;HBA2;RHOC;DUSP6;RHOB;SLC7A5;VWA1;GPRC5C;VAMP5;FOXC1;TAGLN;AHNAK;ADCY4;IFI6;CRIP2;FSTL1;SLC7A1;CRIP1;AKAP2;RELA;FAM129B;FENDRR;FGD5;FOXS1;SAMM50;FRZB;APOL3;NQO1;SLC16A1;PLXDC1;DLC1;CCDC3;AGRN;LIMS2;GIMAP4;PLOD1;GIMAP6;ETS1;GIMAP7;CTGF;CDH6;FOXQ1;CDH5;RGS5;RGS3;SLC39A8;MAP3K6;TGM2;PDGFRB;IGFBP4;LMO2;SLC30A1;EMCN;COL4A2;COL4A1;SLC7A5P2;EOGT;ERG;PHLDB2;KANK3;SNRPB;NOTCH3;NOTCH4;ECSCR;ITPR3;SLC38A11;HIGD1B;SLC9A3R2;MUSTN1;COBLL1;IGFBP7;SLC38A3;BTNL9;GGT5;RBPMS;CAV2;CAV1;ARHGAP29;ISG15;PTPRB;CXCL12;INS-IGF2;RGCC;MOB2;SLCO1A2;DOCK6;ST6GALNAC1;COX4I2;SLC2A1;HSPB1;NRARP;CLEC14A;LAMC1;GPER1;ITGA1;ARAP3;LSR;SHROOM1;DCN;ACTA2;FGR;ADORA2A;COL6A2;OAS3;MMRN2;ITIH5;SLFN5;LEF1;SEMA3G;ATP10A;PRELP;UACA;TTN;DLL4;CLEC3B;CCDC85B;TMEM204;PCDH1;MYO10;LAMB2;NOS3;TIE1;KLF4;KLF2;TBX18;TBX2;EIF3I;NPIPB4;CDK2AP2;TEK |
| **Vascular endothelial cells in Eye** | 49/124 | 3.27129469375754E-07 | 1.7337861876915E-05 | 2.66441151422245 | 39.7874168481916 | ACVRL1;ROBO4;FLT1;TINAGL1;PLXND1;ICAM2;GIMAP4;CLEC14A;GIMAP6;LRRC32;CDH5;SOX18;GRASP;ZNF366;EGFL7;PRKCH;VWF;ARL15;DUSP6;CLDN5;ADORA2A;ERG;SHANK3;ABCG2;KANK3;ENG;RAMP2;ABCB1;EPAS1;NOTCH4;USHBP1;ECSCR;ADCY4;SEMA3G;GATA2;FGD5;PODXL;S1PR1;APOL3;GGT5;PPP1R14A;NOS3;TIE1;PDE2A;ARHGAP29;SMAD7;RGCC;ALPL;TEK |
| **Vascular endothelial cells in Thymus** | 71/234 | 7.56725152743379E-05 | 0.00300798248215493 | 1.7765780364315 | 16.8581187179909 | TINAGL1;FLT1;DOCK9;HIF3A;LDB2;MYLK;NR3C2;PREX2;SHE;MECOM;ZIC2;GJA4;BAALC;DLGAP1;PGM5;KIF1A;RNF152;UBXN6;TGM2;EGFL7;VWF;ITGA1;SOX13;ARAP3;MYRIP;CLDN5;BCAM;EMCN;DAAM2;COL4A1;RHOJ;ITGA6;ERG;SHANK3;KANK3;SHC2;NRN1;EPAS1;USHBP1;ADCY4;ECSCR;TFPI;KALRN;SLC9A3R2;FGD5;ABLIM3;PALMD;PODXL;TSPAN7;SNCG;STOM;A2M;MPDZ;WWTR1;LINC00472;DMTN;MCAM;CAV1;TMEM132B;TIE1;ARHGAP29;TBX2;CDK9;PPFIBP1;PTPRB;MYO1B;ID1;NOSTRIN;TEK;RNF220;RNU4-2 |
| **Vascular endothelial cells in Adrenal** | 40/136 | 0.00467072509070194 | 0.148529057884322 | 1.69311326365876 | 9.08599235731528 | ACVRL1;GPM6A;ROBO4;RAMP2;NRN1;GALNT15;NOTCH4;USHBP1;ECSCR;ADCY4;CLEC14A;LDB2;CX3CL1;EFNB2;SLC9A3R2;DLL4;CDH5;SHE;SOX17;SOX18;PODXL;HYAL2;CA4;CD34;BTNL9;EDN1;TIE1;PDE2A;ARHGAP29;MYRIP;CP;CLDN5;PTPRB;ID1;MMRN2;TEK;SHANK3;VAMP5;KANK3;ENG |
| **Vascular endothelial cells in Placenta** | 28/105 | 0.0538685475707844 | 0.999996079344371 | 1.47487582303338 | 4.30841979717671 | ROBO4;RAMP2;NOTCH4;USHBP1;ECSCR;CLEC14A;NRARP;LDB2;AFF3;LRP6;DLL4;CDH5;SHE;HEY1;SOX18;S1PR1;CD34;GRASP;CGNL1;EGFL7;TIE1;KLF2;CLDN5;RHOJ;EXOC6;ERG;TEK;SHANK3 |

Gene names extracted from enriched terms determined by enrichment analysis using “Descartes Cell Types and Tissue 2021” database.

**Supplementary Table 5: Returned enrichment terms (analysis of mass spectrometry data)**

| **Term** | **Overlap** | **P-value** | **Adjusted P-value** | **Odds Ratio** | **Combined Score** | **Genes** |
| --- | --- | --- | --- | --- | --- | --- |
| **Vascular endothelial cells in Cerebellum** | 84/597 | 3.77E-17 | 4.68E-15 | 3.187628 | 120.543823 | COL18A1;SYNM;TINAGL1;SPARC;TFRC;SLC2A1;HBB;LAMC1;CDH6;CDH5;CSRP1;CA2;BSG;TNKS1BP1;SLC12A7;LMCD1;CD34;CGNL1;TGM2;PDGFRB;EGFL7;ANXA1;VWF;ITGA1;BGN;SLC6A13;SLC6A12;LSR;ACTN4;RHOB;CLDN5;SLC7A5;BCAM;INF2;COL4A2;DAAM2;COL6A2;MMRN2;TAGLN2;MYL9;SHANK3;VAMP3;ENG;KANK3;ABCG2;ITIH5;NOTCH3;TAGLN;ABCB1;CFH;AHNAK;CRIP2;SLC3A2;IQGAP1;SLC7A1;UACA;SLC9A3R2;CLEC3B;RRAS;PALMD;PODXL;CALD1;COBLL1;SERPINH1;IGFBP7;SLC38A5;S100A10;GGT5;SLC16A1;CAV2;LAMB2;NOS3;MCAM;CAV1;FN1;ISG15;PLXDC1;EHD2;MYO1B;SMTN;MYO1C;ESAM;AGRN;ITM2A |
| **Vascular endothelial cells in Cerebrum** | 69/536 | 2.10E-12 | 1.30E-10 | 2.83857992 | 76.3263444 | COL18A1;SYNM;TINAGL1;SPARC;TFRC;SLC2A1;HBB;HSPB1;CLEC14A;LAMC1;CDH6;CDH5;LGALS1;TNKS1BP1;SLC12A7;LMCD1;CD34;CGNL1;TGM2;PDGFRB;ANXA1;ITGA1;BGN;SLC6A13;SLC6A12;LSR;ACTN4;DCN;RHOB;SLC7A5;BCAM;COL4A2;COL6A2;MMRN2;KANK3;ITIH5;NOTCH3;TAGLN;ABCB1;CFH;AHNAK;CRIP2;SLC7A1;UACA;RELA;SLC9A3R2;CLEC3B;RRAS;COBLL1;SERPINH1;IGFBP7;PCDH1;S100A10;GGT5;SLC16A1;CAV2;LAMB2;NOS3;MCAM;CAV1;FN1;ISG15;PLXDC1;EHD2;RPS28;COL1A2;AGRN;ITM2A;LIMS2 |
| **Vascular endothelial cells in Thymus** | 25/234 | 0.0004715 | 0.0194887 | 2.22830141 | 17.0678743 | TINAGL1;DOCK9;MYLK;SLC9A3R2;PALMD;PODXL;TSPAN7;SNCG;STOM;A2M;TGM2;EGFL7;VWF;MCAM;CAV1;ITGA1;SYN2;CLDN5;BCAM;PPFIBP1;MYO1B;DAAM2;ITGA6;SHANK3;KANK3 |
| **Vascular endothelial cells in Eye** | 15/124 | 0.00182175 | 0.05647423 | 2.55189305 | 16.0972341 | GGT5;EGFL7;ABCB1;TINAGL1;VWF;NOS3;CLEC14A;CDH5;CLDN5;PODXL;ALPL;SHANK3;ABCG2;KANK3;ENG |
| **Vascular endothelial cells in Stomach** | 6/36 | 0.00959499 | 0.23628096 | 3.69161793 | 17.1531547 | CLEC3B;CLEC14A;ESAM;A2M;CD34;KANK3 |
| **Stromal cells in Thymus** | 24/282 | 0.01143295 | 0.23628096 | 1.7266519 | 7.72030225 | PDGFRB;LGALS3BP;IGSF8;CBR1;SPARC;SERPINE2;LRP1;ATL1;BGN;PLAT;SORBS3;DCN;C3;UCHL1;PCSK1N;COL1A2;COL5A3;COL6A1;MGP;S100A6;NCAM1;CTNNA2;MFGE8;PFKM |
| **Schwann cells in Adrenal** | 13/132 | 0.01894958 | 0.33567833 | 2.02074039 | 8.01420253 | COL18A1;CNP;HEPACAM;TTYH1;MCAM;ATP1A2;CDH6;BCAN;PTPRZ1;GPC1;PLP1;AGRN;GPM6B |
| **Vascular endothelial cells in Adrenal** | 13/136 | 0.02362679 | 0.36621517 | 1.95461037 | 7.32074734 | GPM6A;CLEC14A;SLC9A3R2;CDH5;CLDN5;PODXL;CA4;MMRN2;CDH13;CD34;SHANK3;KANK3;ENG |
| **Smooth muscle cells in Intestine** | 7/57 | 0.02666625 | 0.36740172 | 2.58392195 | 9.36505407 | TAGLN;FLNA;DSTN;MYH11;MYL9;PDLIM7;MYLK |
| **ENS neurons in Intestine** | 14/157 | 0.03269936 | 0.40547201 | 1.81041944 | 6.19235848 | SYT1;ATP1A3;TUBB4A;UCHL1;GAP43;TUBB2A;MAP1B;EEF1A2;CNTN1;NEFL;NEFM;TAGLN3;MAPT;CTNNA2 |
| **ENS glia in Stomach** | 7/66 | 0.05303433 | 0.55282906 | 2.18872261 | 6.42787527 | PTPRZ1;COL5A3;PLP1;ATP1A2;CRYM;CRYAB;AIF1L |
| **Smooth muscle cells in Heart** | 5/40 | 0.05349959 | 0.55282906 | 2.6336069 | 7.71141524 | PDGFRB;NOTCH3;TAGLN;ARHGEF17;MYH11 |

Gene names extracted from enriched terms determined by enrichment analysis using “Descartes Cell Types and Tissue 2021” database. Genes corresponding to proteins detected during LC-MS/MS were used as input.

**Supplementary Fig. 1: Overlap between top 10% of highly expressed genes from RNA sequencing dataset and dataset from Yang et al. (2022)**

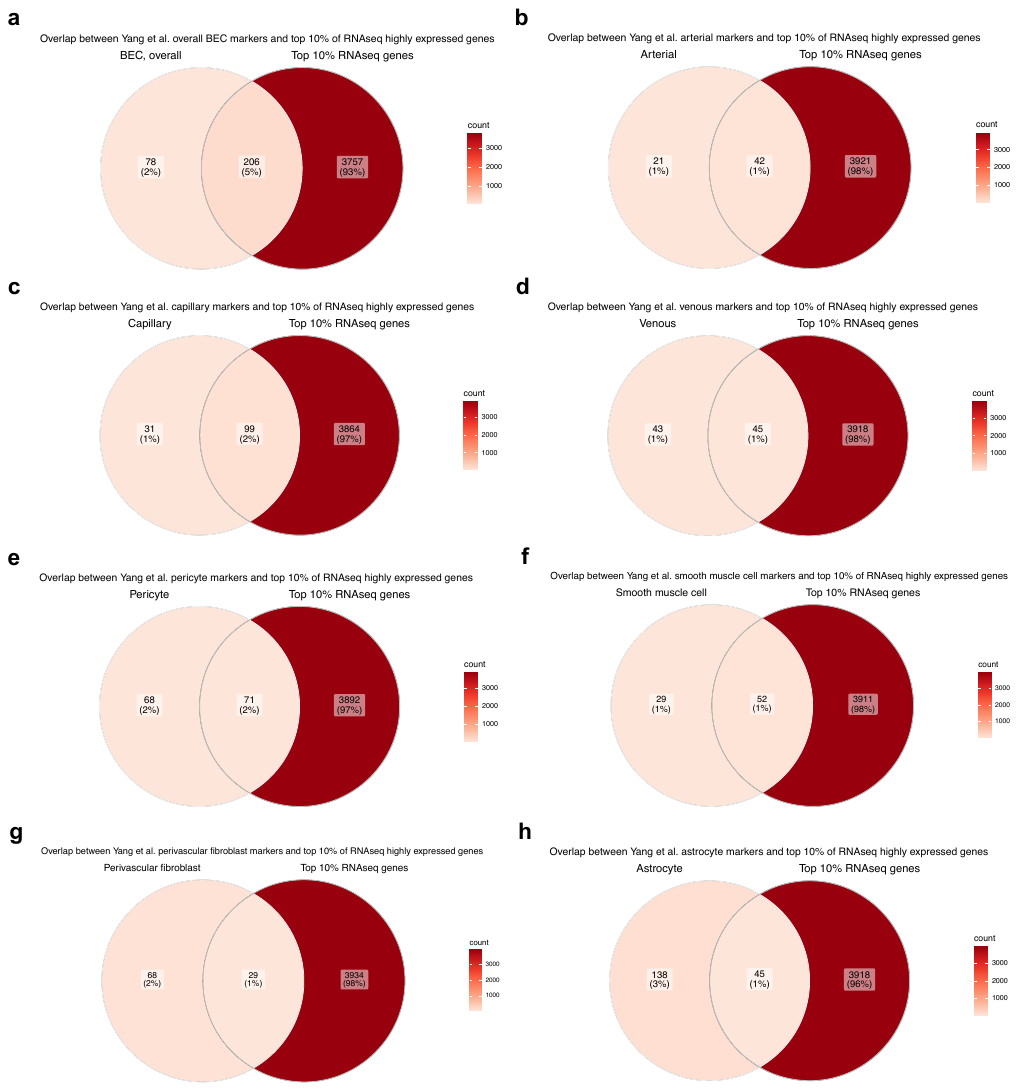

a-h) Validated neurovascular cell-type markers were obtained from Yang et al. (2022) postmortem single-nuclei sequencing dataset (found under supplementary table 2) and compared for potential overlap with the top 10% of most highly expressed genes from our sequencing data.

**Supplementary Fig. 2: LC-MS/MS protocol provides very good coverage of microvessel preparations**

**a)**

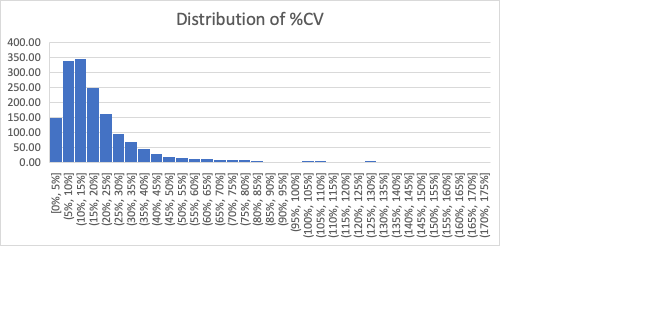

**b)**

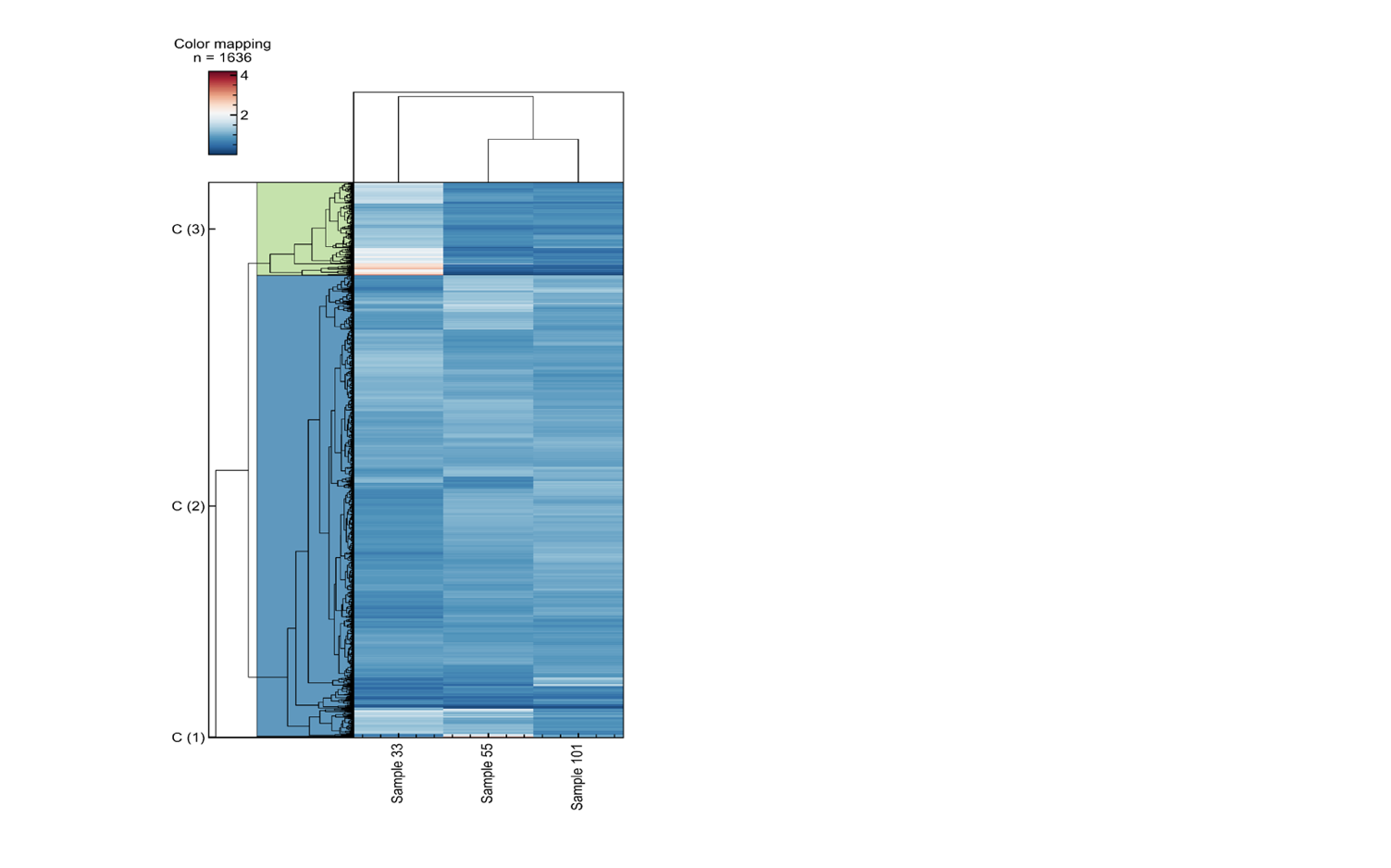

a) Histogram showing the distribution of variation seen for the quantitation of each protein, showing a median of approximately 15%. The coefficient of variation (CV) is calculated by dividing the standard deviation of peptide profiles by the mean, reported as a percentage. b) Heatmap based on tandem mass tag quantitation.
